# Screening the Medicines for Malaria Venture Pathogen Box for invasion and egress inhibitors of the blood stage of *Plasmodium falciparum* reveals several inhibitory compounds

**DOI:** 10.1101/768648

**Authors:** Madeline G. Dans, Greta E. Weiss, Danny W. Wilson, Brad E. Sleebs, Brendan S. Crabb, Tania F. de Koning-Ward, Paul R. Gilson

**Author notes:** Corresponding authors: Madeline G. Dans, Paul R. Gilson.

## Abstract

To identify potential inhibitors of egress and invasion in the asexual blood stage of *Plasmodium falciparum*, we screened the Medicines for Malaria Venture (MMV) Pathogen Box. This compound library comprises of 400 drugs against neglected tropical diseases, including 125 with antimalarial activity. For this screen, we utilised transgenic parasites expressing a bioluminescent reporter, Nanoluciferase (Nluc), to measure inhibition of parasite egress and invasion in the presence of the Pathogen Box compounds. At a concentration of 2 µM, we found 15 compounds that inhibited parasite egress by >40% and 24 invasion-specific compounds that inhibited invasion by >90%. We further characterised 11 of these inhibitors through cell-based assays and live cell microscopy and found two compounds that inhibited merozoite maturation in schizonts, one compound that inhibited merozoite egress, one compound that directly inhibited parasite invasion and one compound that slowed down invasion and arrested ring formation. The remaining compounds were general growth inhibitors that acted during the egress and invasion phase of the cell cycle. We found the sulfonylpiperazine, MMV020291, to be the most invasion-specific inhibitor, blocking successful merozoite internalisation within human RBCs and having no substantial effect on other stages of the cell cycle. This has greater implications for the possible development of an invasion-specific inhibitor as an antimalarial in a combination based therapy, in addition to being a useful tool for studying the biology of the invading parasite.

**Importance:** *Plasmodium falciparum* causes the most severe form of malaria and with emerging resistance to frontline treatments, there is the need to identify new drug targets in the parasite. One of the most critical processes during the asexual blood stage in the parasite’s lifecycle is the egress from old red blood cells (RBCs) and subsequent invasion of new RBCs. Many unique parasite ligands, receptors and enzymes are employed during egress and invasion that are essential for parasite proliferation and survival, therefore making these processes druggable targets. Identifying novel compounds that inhibit these essential processes would further their development into possible antimalarials that would be highly effective at killing asexual RBC stage parasites when used in combination with drugs that target the intraerythrocytic growth phase. These compounds potentially may also be used as novel tools to study the complex biology of parasites to gain further insight into the mechanisms behind egress and invasion.

## Introduction

Malaria remains a significant global health burden with an estimated 219 million cases worldwide in 2017, resulting in 435 000 deaths (1). Of the *Plasmodium* species known to infect humans, *Plasmodium falciparum* remains the deadliest and is therefore a focus in the fight to eradicate malaria. There is emerging resistance of *P. falciparum* to the gold-standard artemisinin combination therapies (ACTs) whereby delayed parasite clearance has been observed in regions of Southeast Asia (1–4). With the spread of resistance to artemisinin and its partner drugs, it is vital that novel therapeutics are developed and ready to deploy when ACTs are rendered ineffective.

The red blood cell (RBC) stage of infection causes the clinical symptoms of malaria. Asexual blood stage parasites progress through a series of developmental phases that begin with the intracellular ring stage, followed by the trophozoite stage and concludes with the DNA-replicative schizont phase. From mature schizonts, approximately 20 invasive merozoites emerge from the nutrient-deprived infected RBC (iRBC) after the breakdown of the parasitophorous vacuole membrane (PVM), followed by the rupture of the RBC membrane (5, 6). These merozoites rapidly invade new RBCs and it has been shown that these processes require a temporal and spatial cascade of signalling kinases and proteases (7–9). Compounds that inhibit these enzymes such as the cGMP dependent protein kinase G (PKG) specific inhibitor, Compound 1 (C1), have proven to be valuable tools in helping to decipher the roles of various egress and invasion proteins (10, 11).

Following egress, the merozoite secretes proteins from its unique secretory organelles; the rhoptries, micronemes, and dense granules, which enable the merozoite to invade new RBCs in a complex multi-step process that is still not fully understood (12–14). The primary contact between a merozoite and a RBC occurs via a multi-protein complex containing merozoite surface protein 1 (MSP1) (15, 16) and possibly heparin sulfate proteoglycan receptors on the RBC surface, since heparin inhibits this interaction (17, 18). Subsequent stronger attachment between merozoites and RBCs is through the binding of merozoite proteins, namely, reticulocyte binding like homologs (Rhs) and erythrocyte binding proteins (EBAs), which are secreted from the rhoptries and micronemes, respectively (19–22). Downstream binding of Rh5 to the RBC receptor, basigin, then possibly activates the secretion of the rhoptry neck protein complex (RON) which becomes embedded into the RBC surface (23). Here, RON2 interacts with apical membrane antigen 1 (AMA1) which is secreted from the micronemes onto the merozoite surface, leading to the formation of a tight junction between the invading merozoite and RBC (24, 25). The parasite’s actin-myosin motor applies a penetrative force to propel the merozoite into the RBC, during which the merozoite envelops itself in the RBC membrane, forming the PVM (26, 27). Less than a minute after invasion, the RBC undergoes echinocytosis, a morphological change from its normal biconcave shape to a stellate form, hypothesised to be caused by an efflux of ions or a disruption to the phospholipid bilayer of the RBC upon secretion of rhoptry proteins (23, 28, 29). Since many of the protein-protein interactions, signalling cascades and enzymes required for egress and invasion are unique to parasites, they could represent novel drug targets that may be effective antimalarials when used in combination with drugs that act during the intraerythrocytic stage (reviewed in (30)).

To facilitate open source drug discovery for antimalarials, Medicines for Malaria Venture (MMV) has released a series of small compound libraries, screened for their ability to inhibit the RBC stage of the parasite’s lifecycle (http://www.mmv.org/research-development). The first compound library, termed the Malaria Box (31), was phenotypically screened by Subramanian *et al.* (2018) for blood stage egress and invasion inhibitors which resulted in the identification of 26 compounds that inhibited schizont to ring transition by greater than 50% (32). The second-generation library released by MMV was labelled the Pathogen Box, and it contains 400 compounds against neglected tropical diseases, 125 from the Malaria disease set and 15 with activity against the related parasite, *Toxoplasma gondii*. We have screened the Pathogen Box using a bioluminescent semi-high throughput system to identify inhibitors of RBC egress and invasion, identifying 15 and 24 inhibitors, respectively with these properties. After removing compounds with known, non-invasion related targets or those that did not inhibit parasite growth, we performed a detailed analysis of 11 of these compounds by studying their effects upon egress and invasion.

## Results

### Development and validation of bioluminescent egress and invasion screening assay

To screen the MMV Pathogen Box for egress and invasion inhibitors, we used *P. falciparum* 3D7 parasites expressing a Nanoluciferase (Nluc) that is exported into the cytoplasmic compartment of the iRBC (previously described in (33)) in an assay we termed the Nluc invasion assay (Figure 1A). Briefly, in this assay late stage schizonts were purified and added to RBCs in the presence of inhibitors for four hours. The culture media was then collected to measure bioluminescence in relative light units (RLU) of the Nluc released upon schizont rupture. The iRBCs were then treated with 5% isotonic sorbitol to lyse remaining schizonts, leaving intact newly invaded ring stage parasites. Since it was not possible to measure Nluc in the new ring stage parasites as Nluc expression was too low (33), the iRBCs were grown for ∼24 hours until trophozoites (parasite age range 24-28 hours post invasion (hpi)). The trophozoite iRBCs were then lysed and degree of invasion was inferred by RLU levels.

**Figure 1.**
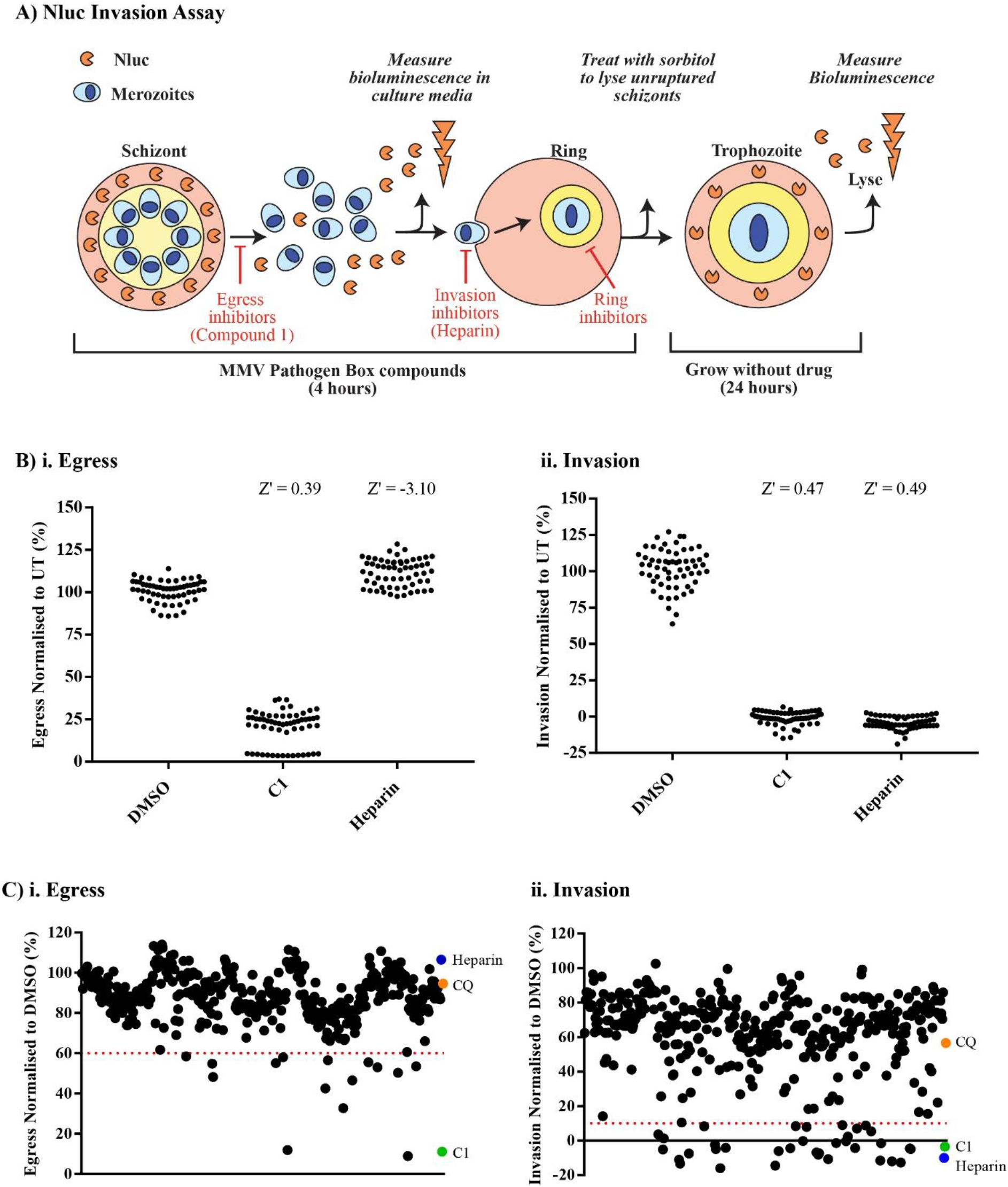
Blood stage parasites expressing an exported nanoluciferase reporter protein enables the quantification of egress and invasion and identified inhibitors of these processes in the MMV Pathogen Box. **A)** Schematic of the nanoluciferase (Nluc) invasion assay set-up whereby purified late schizonts expressing an exported Nluc were treated with compounds for 4 hours. Egress inhibitors, such as Compound 1 (C1), prevent release of Nluc which was detected when the bioluminescence of the growth media was measured after 4 hours incubation. The infected RBCs were then treated with 5% sorbitol to lyse schizonts, leaving newly infected ring-stage parasites behind. The ring-stage parasites were grown until trophozoites when they were lysed and their total Nluc levels measured to infer the degree of invasion. The invasion inhibitor, heparin, was used as a control compound. B) Egress (i) and invasion (ii) of Nluc parasites following a 4 hour treatment with compound vehicle DMSO, C1 or heparin. (i) The Z score of 0.39 for C1 demonstrates an acceptable separation band between positive and negative controls. (ii) For parasite invasion, both C1 and heparin were used as positive controls which was reflected in their Z scores of 0.47 and 0.49, respectively. Values have been normalised and expressed as a percentage of untreated (UT) parasites. Sixty replicates were performed for each condition over 3 independent experiments. Z’ indicates Z score. C) Using the Nluc egress/invasion assay, the Pathogen Box compounds were screened at 2 µM and it was found that 15 compounds reduced egress rate <60% (i) and 36 compounds reduced invasion rate <10% (ii). All values have been normalised to compound vehicle DMSO. Each dot represents the mean of a compound from 3 biological replicates. Dotted lines indicate cut-off values of 60% and 10% for egress and invasion inhibition, respectively. Positive control compounds, heparin and C1, were used at 100 µg/mL and 4 µM, respectively. Chloroquine (CQ) was included as a negative control at 75 nM and DMSO vehicle control was used at 0.02%.

To validate this assay’s suitability for high throughput screening (HTS), we generated Z scores using the well-characterised egress and invasion inhibitors, Compound 1 (C1) and heparin, respectively as positive controls for inhibition of egress and invasion, respectively (11, 23, 34–36). For both egress and invasion, the assay achieved acceptable separation bands between its positive (C1 and heparin) and negative drug vehicle DMSO control, producing Z scores for egress and invasion of 0.39 and 0.49, respectively (Figure 1B). C1, heparin and R1 peptide, which blocks the interaction between AMA1 and RON2 (37), were also tested at multiple concentrations in the Nluc invasion assay which produced dose-response curves from which we could derive the half maximal effective concentration (EC_50_) for egress and invasion (Figure S1). It should be noted that egress inhibitors, like C1, also inhibit invasion in this assay because the merozoites are unable to escape the iRBC (Figure 1A).

### Screen of the Pathogen Box compounds reveals many egress and invasion inhibitory compounds

The Pathogen Box compounds were screened at 2 µM for invasion and egress inhibition using the Nluc invasion assay. Based on control compound activity in the Z score assays, cut-offs for positive hits of egress inhibitors were set at <60% and for invasion inhibitors set at <10%. From the screen, we found 15 compounds that reduced egress to <60% and 36 compounds that reduced invasion to <10% when normalised to DMSO at 100% (Figure 1C and Table S1). The hits from the Nluc screen then underwent a compound triaging process to identify compounds that were to be investigated further (Tables 1 and 2). Of the 15 compounds that inhibited egress, MMV688274 was removed since a counter screen performed with parasite lysate indicated the compound inhibited Nluc activity (Figure S2, Table 1). Due to the Nluc invasion assay’s design, inhibitors of early ring-stage parasites (0-4 hpi) could be exposed to the parasites for up to four hours, resulting in false positives for invasion inhibition (Figure 1A). As such, compounds were tested for early ring stage growth inhibition activity and this resulted in the removal of one of the egress hit compounds, MMV667494, leaving a remaining 13 compounds targeting egress (Figure 2A, Table 1). Since we were interested in studying compounds with novel egress targets, we further triaged our list of egress inhibitors to remove compounds with known targets and mechanisms of action. One of these was compound MMV688703 which was found to be structurally identical to C1 and included in the Pathogen Box as it targeted *Toxoplasma gondii* PKG (Table 1) (38). MMV030734 was also excluded since it acted upon *P. falciparum* calcium dependent protein kinase 1 (PfCDPK1), known to be involved in microneme secretion and activation of the actin-myosin motor (Table 1) (39, 40). Six compounds which likely target PfATP4 were also removed since PfATP4 is involved in the efflux of excess Na^+^ from the parasite’s cytoplasm and is therefore unlikely to be directly involved in egress or invasion which we confirmed using a purified merozoite invasion assay with the known PfATP4 inhibitor, cipargamin (Table 1, Figure S3B) (41, 42). MMV016838 was found to be the parent compound of an antimalarial in clinical development, M5717, that targets elongation factor 2 (PfEF2) (43, 44). However, as MMV016838 did not inhibit ring stage parasites like other PfEF2 inhibitors in the Pathogen Box that were hits in our screen (MMV667494 and MMV634140 (45)), this compound was not removed. Two additional compounds from the tuberculosis disease set were also excluded because they exhibited no antimalarial activity in the *P. falciparum* blood stage that had been previously performed by Duffy*, et al*. (2017) (Table 1) (46). This left three compounds, MMV011765, MMV016838 and MMV019993 of which we were able to obtain additional quantities of the first two compounds from commercial sources and MMV to further study.

**Figure 2.**
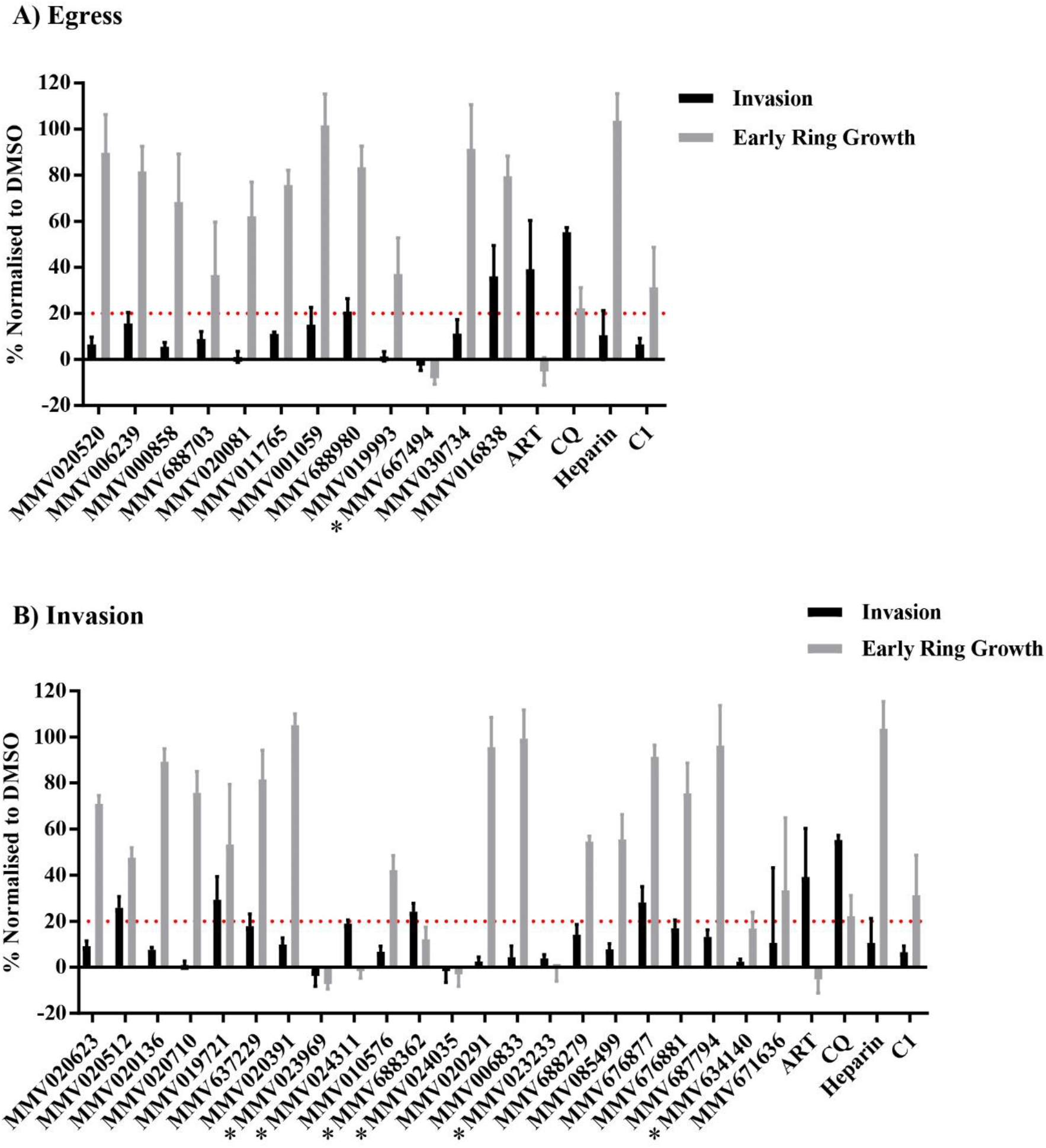
Multiple egress and invasion inhibitors also target the early ring-stage of intracellular growth. The lead compounds from the egress screen **(A)** and the invasion screen **(B)** were assessed for their ability to inhibit early ring parasites (4-8 hours post invasion) at a concentration of 2 µM in the Nluc invasion assay. One compound from the egress screen and 6 compounds from the invasion screen potently inhibited ring stage parasites (*). Artemisinin (ART) was included as a positive control for ring stage inhibition at a concentration of 25 nM, and negative controls were chloroquine (CQ), compound 1 (C1) and heparin which were used at concentrations of 75 nM, 4 µM and 100 µg/mL, respectively. Values were normalised to DMSO at a concentration of 0.02%. Dotted line indicates 20% cut-off for early ring stage inhibitors. Error bars represent standard deviation of 3 biological replicates.

**Table 1.**
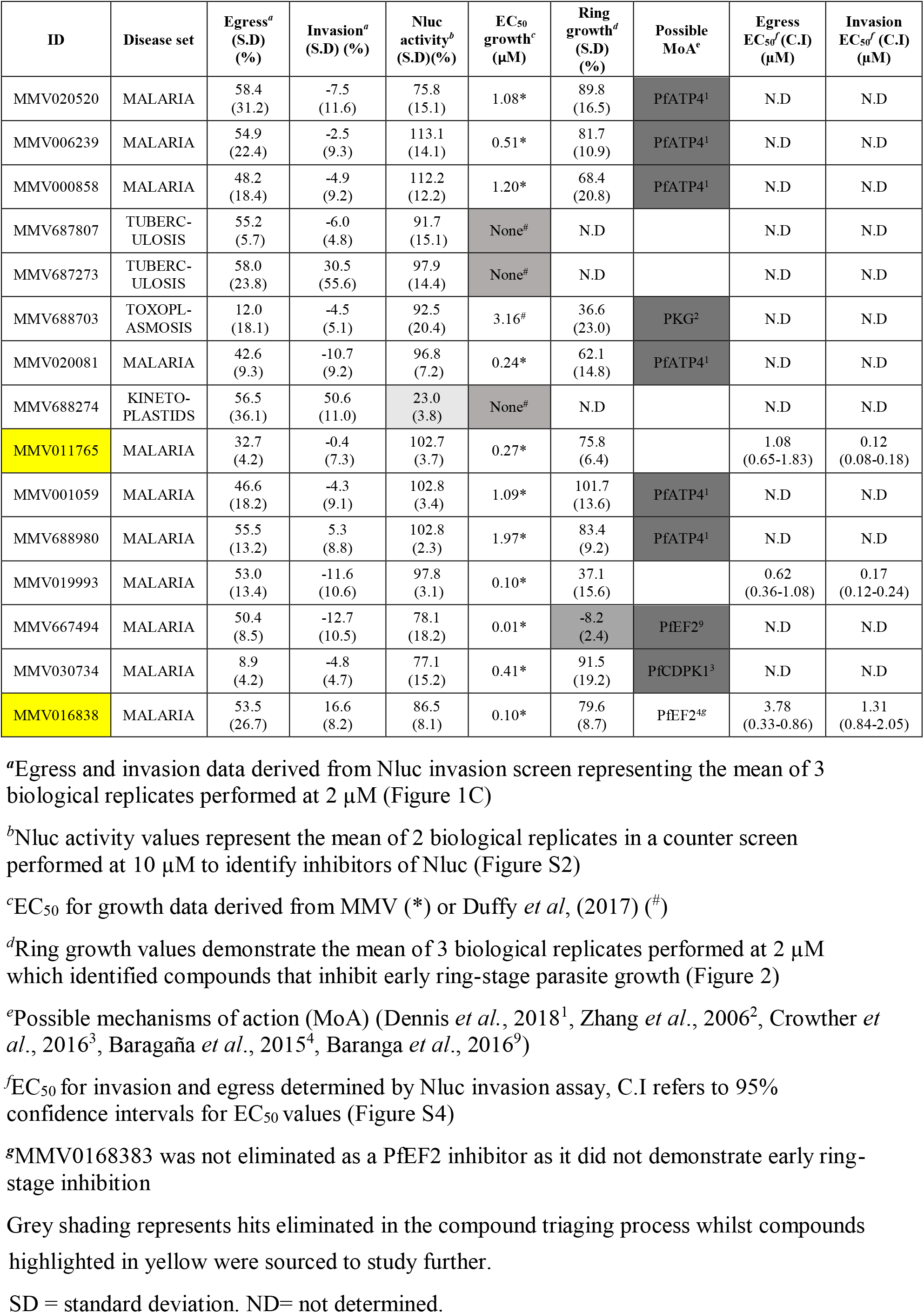
15 Pathogen Box compounds identified as egress hits.

Of the 36 compounds which reduced invasion to <10%, 12 of these were also inhibitors of egress and were subsequently removed, leaving a remaining 24 invasion-specific inhibitory compounds (Table 2). Two reference compounds (mefloquine (MMV000016) and pentamidine (MMV000062)) were also excluded. The invasion hits were also tested for their early ring stage growth inhibition activity and this resulted in the removal of six compounds (MMV023969, MMV024311, MMV688362, MMV024035, MMV023233, MMV634140) (asterisks, Figure 2B and Table 2). Compounds targeting PI4K, the mitochondrial cytochrome bc1 complex, DNA machinery, PfEF2 and PfATP4 were removed since their proteins targets have likely non-invasion related roles (Table 2) (41, 45–50). This left nine compounds targeting invasion which we were able to obtain supplementary amounts of from commercial sources or MMV to conduct phenotypic analyses.

**Table 2.** 24 Pathogen Box compounds identified as invasion-specific hits.

| ID | Disease set | Invasion<sup>a</sup><br>(S.D) (%) | Nluc activity<sup>b</sup><br>(S.D) (%) | EC<sub>50</sub><br>growth<sup>c</sup><br>(μM) | Ring<br>growth<sup>d</sup><br>(S.D) (%) | Possible<br>MoA<sup>e</sup> | Invasion EC<sub>50</sub><sup>f</sup><br>(C.I) (μM) |
| --- | --- | --- | --- | --- | --- | --- | --- |
| MMV000062 | REF.<br>(PENTAMIDINE) | 3.6 (10.6) | 96.4 (6.1) | N/A | N.D |  | N.D |
| MMV020623 | MALARIA | -5.1 (11.4) | 90.1 (6.9) | 0.94* | 71.0 (3.6) | PfATP4<sup>1</sup> | N.D |
| MMV020512 | MALARIA | 1.2 (10.8) | 94.1 (8.4) | 0.48* | 47.6 (4.3) |  | 0.86<br>(0.37-1.98) |
| MMV020136 | MALARIA | -11.1 (10.6) | 91.8 (30.8) | 1.10* | 89.3 (5.6) | PfATP4<sup>1</sup> | N.D |
| MMV020710 | MALARIA | -13.3 (10.2) | 81.9 (22.6) | 0.24* | 75.8 (9.2) | PfATP4<sup>1</sup> | N.D |
| MMV019721 | MALARIA | 10.6 (13.3) | 90.9 (22.1) | 0.50* | 53.3 (26.1) |  | 1.78<br>(1.01-3.15) |
| MMV637229 | TRICHURIASIS | 8.3 (9.5) | 105.7 (17.3) | 2.20<sup>#</sup> | 81.6 (12.7) |  | 0.50<br>(0.33-0.74) |
| MMV000016 | REF.<br>(MEFLOQUINE) | -15.9 (11.1) | 105.5 (16.3) | N/A | N.D |  | N.D |
| MMV020391 | MALARIA | -4.2 (11.7) | 101.9 (19.6) | 0.93* | 105.1 (4.9) | PfATP4<sup>1</sup> | N.D |
| MMV023969 | TUBERCULOSIS | -14.5 (8.7) | 100.2 (10.2) | 0.59<sup>#</sup> | -7.2 (2.2) |  | N.D |
| MMV024311 | TUBERCULOSIS | 8.4 (14.4) | 104.1 (20.6) | 0.75<sup>#</sup> | -1.6 (3.2) |  | N.D |
| MMV010576 | MALARIA | -0.3 (13.7) | 93.8 (5.1) | 0.10* | 42.3 (6.2) | PI4K<sup>5,6</sup> | N.D |
| MMV688362 | KINETOPLASTIDS | 7.9 (10.9) | 98.1 (5.7) | 0.59<sup>#</sup> | 12.2 (5.3) | DNA binding<br>agent<sup>8,10</sup> | N.D |
| MMV024035 | MALARIA | -6.8 (9.6) | 87.4 (15.2) | 0.57* | -3.0 (5.2) |  | N.D |
| MMV020291 | MALARIA | -8.2 (9.2) | 100.6 (8.8) | 0.91* | 95.6 (12.9) |  | 0.43<br>(0.25-0.75) |
| MMV006833 | MALARIA | -7.3 (7.2) | 95.3 (12.4) | 0.30* | 99.3 (12.5) |  | 0.17<br>(0.10-0.27) |
| MMV023233 | MALARIA | -1.4 (4.6) | 104.5 (0.01) | 0.16* | -0.7 (5.4) |  | N.D |
| MMV688279 | KINETOPLASTIDS | 9.0 (11.2) | 88.8 (19.5) | 0.32<sup>#</sup> | 54.5 (2.4) |  | 0.43<br>(0.20-0.92) |
| MMV085499 | MALARIA | 2.3 (10.5) | 90.1 (4.0) | 0.10* | 55.5 (10.8) | PI4K<sup>7</sup> | N.D |
| MMV676877 | MALARIA | 7.1 (4.6) | 91.2 (1.0) | 0.66* | 91.4 (5.1) |  | 1.36<br>(0.68-2.85) |
| MMV676881<sup>T</sup> | MALARIA | 8.8 (5.9) | 98.4 (4.7) | 0.90* | 75.6 (13.1) | Cysteine<br>proteases<sup>11,12</sup> | 0.17<br>(0.09-0.31) |
| MMV687794 | MALARIA | -1.5 (9.7) | 86.9 (2.6) | 0.01* | 96.2 (17.4) |  | 0.31<br>(0.16-0.60) |
| MMV671636 | ONCHOCERCIASIS | -12.0 (8.6) | 90.3 (20.2) | 1.01<sup>#</sup> | 33.4 (31.5) | Mitochondria<sup>8</sup> | N.D |
| MMV634140 | MALARIA | -4.6 (10.9) | 90.1 (10.6) | 0.10* | 16.8 (7.05) | PfEF2<sup>9</sup> | N.D |

<sup>a</sup>Invasion data derived from Nluc invasion screen representing the mean of 3 biological replicates performed at 2 μM (Figure 1C)
<sup>b</sup>Nluc activity values represent the mean of 2 biological replicates performed at 10 μM in a counter screen to identify inhibitors of Nluc (Figure S2)
<sup>c</sup>EC<sub>50</sub> for growth data derived from MMV (\*) or Duffy *et al*, (2017) (#)
<sup>d</sup>Ring growth values demonstrate the mean of 3 biological replicates performed at 2 μM which identified compounds that inhibit early ring parasite growth (Figure 2)
<sup>e</sup>Possible mechanisms of action (MoA) (Dennis *et al.*, 2018<sup>1</sup>, Younis *et al.*, 2012<sup>5</sup>, Paquet *et al.*, 2017<sup>6</sup>, Le Manach *et al.*, 2016<sup>7</sup>, Duffy *et al.*, 2017<sup>8</sup>, Baranga *et al.*, 2016<sup>9</sup>, Rodríguez *et al.*, 2008<sup>10</sup>, Mott *et al.*, 2011<sup>11</sup>, Veale, 2019<sup>12</sup>)
<sup>f</sup>EC<sub>50</sub> for invasion determined by Nluc invasion assay, C.I refers to 95% confidence intervals for EC<sub>50</sub> values (Figure S4)
<sup>T</sup>MMV676881 identified in the screen as invasion inhibitor was found to be an egress inhibitor upon further characterisation
Grey shading represents hits eliminated in the compound triaging process whilst compounds highlighted in yellow were sourced to study further
SD = standard deviation. ND= not determined. N/A= not applicable.

### Categorising egress and invasion inhibitory compounds

To identify the phenotypes of these invasion and egress inhibitors, Giemsa-stained blood smears were examined from schizonts that had been treated for four hours with 2 µM of the hit compounds (Figure 3). It was found that the egress inhibitor, MMV011765, halted schizont maturation at the point when the merozoites had formed but had not yet physically separated possibly due to an intact PVM (Figure 3A). The egress inhibitor, MMV016838 did not appear to inhibit egress or invasion at 2 µM when compared to vehicle control. Therefore, the concentration was increased to 20 µM (10 x EC_50_ of egress, Figure S4A.i) upon which a similar inhibition of schizont/merozoite maturation phenotype to MMV011765 and C1 was observed (Figure 3A). Based on the Nluc screen, MMV676881 was predicted to be an invasion inhibitor, however Giemsa stained smears indicated that MMV676881 prevented schizont egress, probably after breakdown of the PVM but not the RBC membrane since the merozoites appeared physically separated within the schizont (Figure 3B).

**Figure 3.**
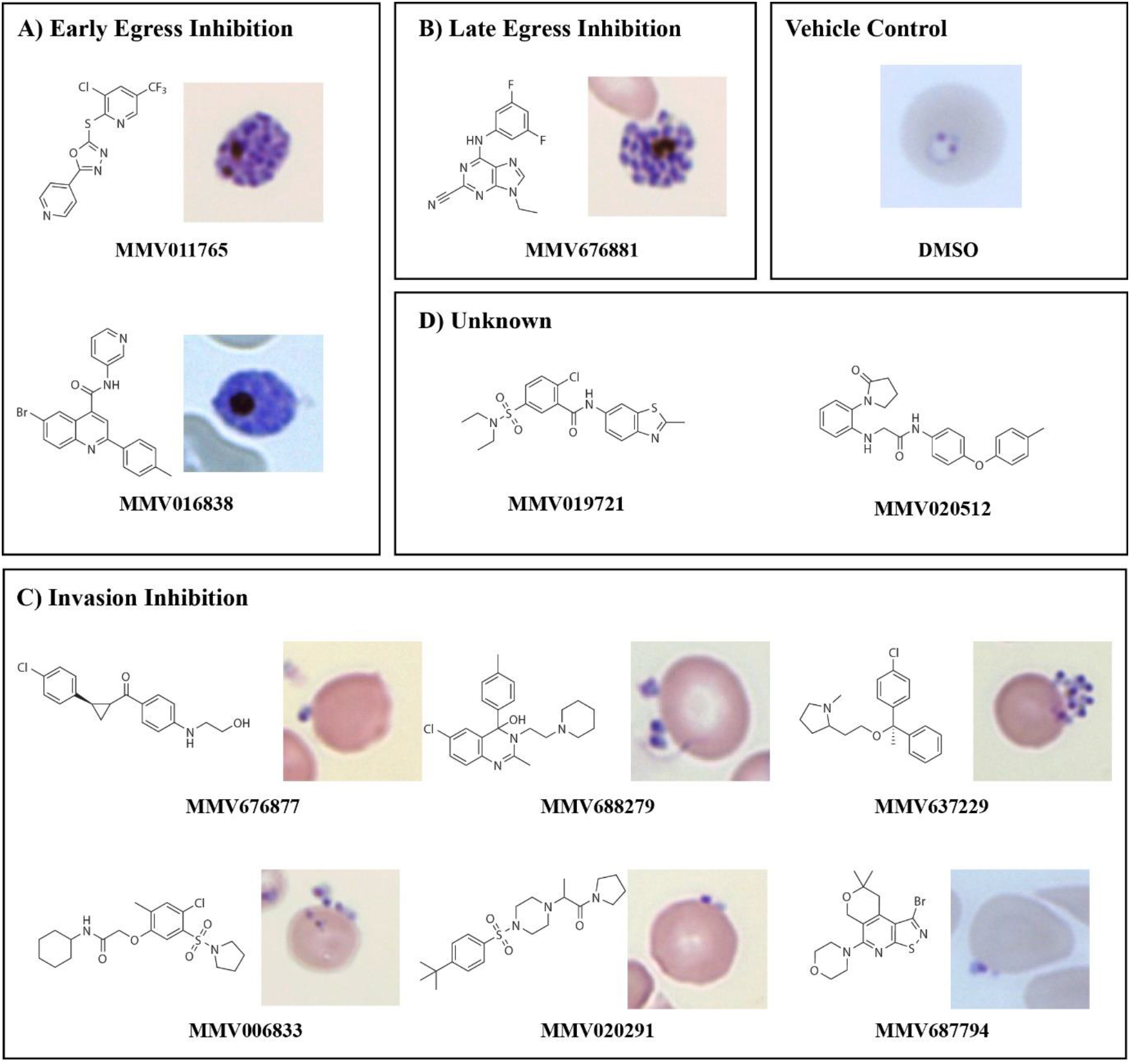
Giemsa stained thin blood smears of the lead compounds for egress and invasion inhibition depict two early egress inhibitory compounds, one late egress inhibitor and six invasion blockers. Giemsa-stained smears following a 4 hour treatment of schizonts with 2 µM of Pathogen Box compounds (or 20 µM of MMV016838) were used to categorise the lead compounds into either early egress inhibitors **(A)**, late egress inhibitors **(B),** invasion inhibitors **(C)** or unknown **(D)** 0.1% DMSO depicts vehicle control.

Of the remaining eight invasion inhibitors identified in the screen, six (MMV676877, MMV006833, MMV637229, MMV020291, MMV688279, MMV687794) demonstrated a degree of invasion inhibitory activity by Giemsa smears since less rings were observed than in the DMSO vehicle control. In addition, merozoites were observed to be either stuck on the outside of the RBCs or not properly differentiated into ring-stage parasites (Figure 3C). The remaining two compounds identified as invasion inhibitors from the screen (MMV019721 and MMV020512) did not appear to have inhibitory effects on egress or invasion at 2 µM when compared with the vehicle control by Giemsa smears and therefore the concentration was increased to approximately 10 x EC_50_ of invasion (10 µM and 6 µM, respectively, Figure S4B). Whilst this reduced the number of new rings compared to the vehicle control, there were no obvious phenotypes to explain how the compounds were inhibiting the invasion process (Figure 3D).

### Egress inhibitors function at two stages of schizont maturation and do not affect merozoite invasion of RBCs

To further characterise the egress inhibitors, live schizonts were examined under culture conditions by brightfield microscopy after four hours of drug treatment. MMV011765 and MMV016838 treatment revealed they were similar to C1-arrested schizonts where the merozoites were indistinct from each other, possibly because they were spatially confined with intact PVM and RBC compartments (Figure 4A.i). To test if this inhibition was reversible, schizonts expressing Nluc were treated with C1, MMV011765 or MMV016838 for two hours to arrest egress which was confirmed due to reduced Nluc activity in the growth media relative to a DMSO control (Figure 4A.ii). To measure reversibility, the compounds were washed out and egress was allowed to proceed for a further four hours. Nluc activity in the growth media indicated egress had resumed for the reversible inhibitor, C1, and MMV016838 but not for MMV011765, indicating the latter is an irreversible egress inhibitor (Figure 4A. ii).

**Figure 4.**
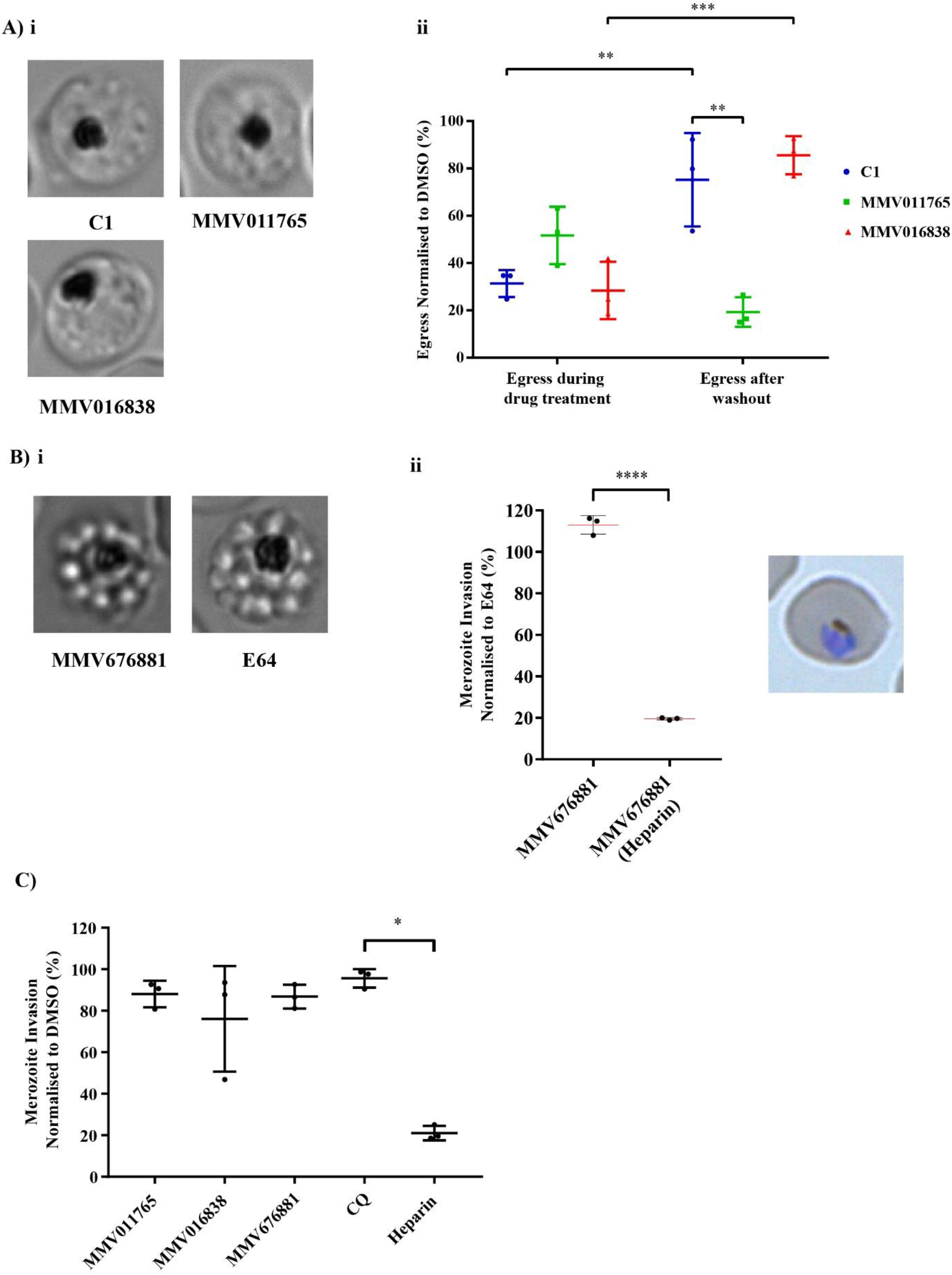
Treatment of schizonts with egress inhibitors reveals some compounds are reversible and/or do not inhibit invasion of purified merozoites. **A)** (i) Brightfield microscopy reveals treatment of schizonts with 10 µM MMV011765 and MMV016838 blocks the maturation and separation of merozoites similar to treatment with 4 µM C1. (ii) Measurement of egress during and after the removal of inhibitory compounds as determined by the release of Nluc into the growth media indicates MMV011765 is an irreversible inhibitor whilst MMV016838 is a reversible inhibitor, like C1.Values have been normalised to 0.1% DMSO with concentrations of MMV compounds at 10 µM and C1 at 4 µM. **B)** (i) Live cell microscopy demonstrates that MMV676881 prevents the release of merozoites, similarly to the protease inhibitor E64 by producing PVM-enclosed merozoite structures. Single frames of videos are shown in (i) (Supplementary video 1, 2) after a 4 hour treatment of schizonts with 10 µM MMV676881 and 10 µM E64. (ii) Mechanical rupture of RBC membranes after 4 hour treatment with 10 µM MMV676881 and 10 µM E64 demonstrated the merozoites were viable and invasion competent with exposure to MMV676881 resulting in a similar degree of invasion as E64-treatment as determined by measurement of Nluc activity 24 hours later. Control compound heparin inhibited merozoite invasion of MMV676881 treated schizonts. A Giemsa stained smear of a trophozoite at 24 hours post invasion after MMV676881 schizont treatment showing that merozoites invaded and progressed to trophozoites normally. All values have been normalised to 10 µM E64. **C)** Merozoite invasion assays demonstrate MMV011765, MMV016838 and MMV676881 at 10 x EC_50_ for growth were specific for egress inhibition with merozoite invasion remaining relatively unaffected. No significant difference between the MMV compounds and chloroquine (CQ) was seen, in contrast to heparin. Values have been normalised to 0.1% DMSO. Concentrations of control compounds, heparin and chloroquine were 100 µg/mL and 75 nM, respectively. Statistical analysis for A.ii was performed via two-way ANOVA; B.ii, unpaired t test; C, one-way ANOVA using GraphPad Prism. * indicates p < 0.05, ** indicates p <0.01, *** indicates p <0.001 and **** indicates p <0.0001. No bar indicates not significant. Error bars represent the standard deviation of 3 biological replicates.

Next, schizonts were treated with the later acting MMV676881 and when visualised by brightfield microscopy, PVM-enclosed merozoite structures (PEMS) were observed, resembling treatment with the cysteine protease inhibitor, E64 (Figure 4B.i, Supplementary Video 1 and 2). It should be noted that E64 does not prevent Nluc release in the Nluc invasion assay, similarly to MMV676881 (Figure S3A). To further support MMV676881 acting as an E64-like inhibitor, a merozoite viability assay was performed whereby purified schizonts were treated with either MMV676881 or E64 to induce PEMS, and mechanically broken open to release merozoites to allow invasion of new RBCs. A negative control containing heparin was also included to block invasion. Ring-stage parasites were grown for 24 hours and the degree of invasion was inferred by measuring the Nluc activity of the whole culture. The Nluc activity of trophozoites treated with MMV676881 at schizonts in the previous cycle was, on average, 108% of those treated with E64, with heparin reducing invasion to 20% (Figure 4B. ii, p<0.0001), demonstrating that both MMV676881 and E64 block merozoite egress but do not affect the merozoites ability to invade RBCs. This strengthens support for MMV676881 acting similarly to an E64-like egress inhibitor by preventing breakdown of RBC membranes, without affecting merozoite viability.

The three egress inhibitors, MMV011765, MMV016838 and MMV676881 were also tested for their ability to specifically inhibit merozoite invasion in invasion assays with purified merozoites (51–53). Briefly, purified schizonts were treated with E64 until the formation of PEMS, then mechanically ruptured to release the merozoites. The merozoites were rapidly added to fresh RBCs in the presence of 10 x EC_50_ of the compounds and left to invade the RBCs at 37°C for 30 minutes. The compounds were then washed out of the new ring stage parasites and cultured for a further 24 hours with quantification of invasion performed by measuring Nluc activity present in trophozoite iRBCs. This revealed that MMV011765, MMV016838 and MMV676881 did not affect merozoite invasion of RBCs, with no differences observed between the egress inhibitors and the antimalarial, chloroquine, which does not inhibit invasion. In contrast, the control compound heparin dramatically reduced the degree of RBC invasion (Figure 4C). Taken together, these findings indicate that MMV011765, MMV016838 and MMV676881 are inhibitors of schizont egress at early and late stages of schizont maturation and do not inhibit merozoite invasion of RBCs.

### Two inhibitors, MMV020291 and MMV006833, appear to specifically block merozoite entry into RBCs

The eight compounds (MMV676877, MMV006833, MMV637229, MMV020291, MMV688279, MMV687794, MMV019721 and MMV020512) that were predicted to inhibit invasion from the Nluc screen (six of which displayed invasion inhibitory effects by morphological examination of Giemsa stained smears) were tested for a direct inhibition of merozoite invasion by performing purified merozoite invasion assays as described above. This revealed that only one of the compounds, MMV020291, could directly block merozoite invasion to a similar degree to that of the heparin control (Figure 5A). MMV637229, MMV006833 and MMV020512 had an intermediate effect, whereas the other compounds, MMV676877, MMV019721, MMV687794 and MMV688279 caused negligible invasion inhibition, similar to the chloroquine (Figure 5A).

**Figure 5.**
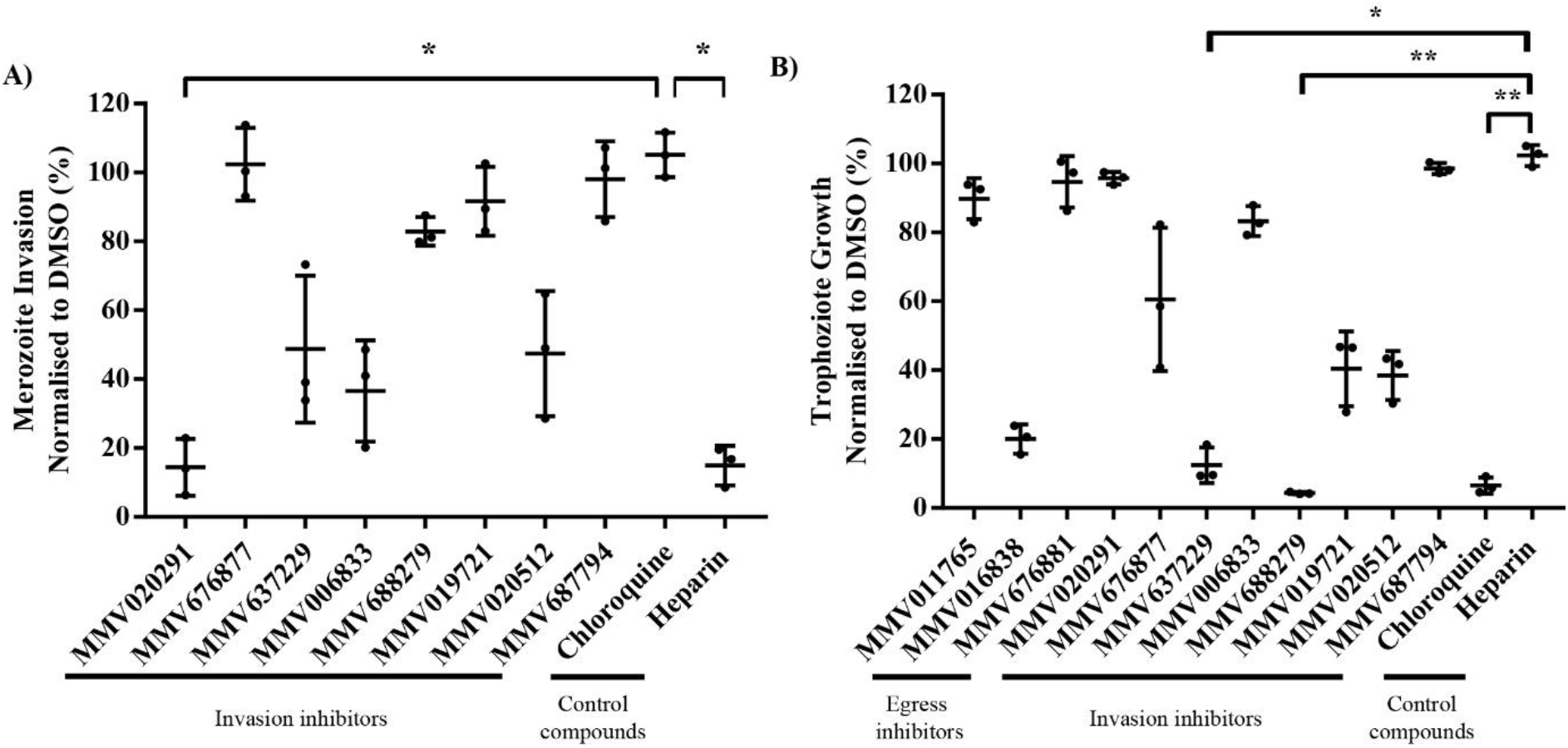
MMV020291 is the most specific invasion inhibitory compound. **A)** Merozoite invasion assays demonstrates 1 of 8 the invasion inhibitors identified from the screen, MMV020291, is the only compound to directly block merozoite invasion into RBCs to a similar degree as heparin. MMV637229, MMV006833 and MMV020512 induce intermediate invasion inhibitory effects whereas the remaining compounds have little effect. **B)** Growth of trophozoite stage parasites after exposure to the lead compounds for 4 hours demonstrates that multiple compounds (MMV688279, MMV637229, MMV016838) reduce trophozoite growth with MMV688279 causing significant growth reduction when compared with control compound, heparin. Pathogen Box compounds were tested at 10 x EC_50_ of growth (or 5 x EC_50_ for MMV637229) whilst control compounds heparin and chloroquine were used at concentrations of 100 µg/mL and 75 nM, respectively with values normalised to 0.1% DMSO. Error bars represent the standard deviation of 3 biological replicates. Statistical analysis performed via one-way ANOVA in GraphPad Prism between chloroquine (A) and heparin (B) and Pathogen Box compounds. *indicates p < 0.05. No bar indicates not significant.

We hypothesised that the intermediate invasion inhibitory compounds may be exerting their invasion inhibitory effects by causing general growth defects during the window of merozoite egress and/or invasion. We ascertained this by measuring the compounds’ effects on other stages in the asexual lifecycle. Whilst activity against ring-stage parasites (4-8 hpi), was a criterion for elimination in compound triaging from the screen, it was possible that these compounds may be active at other stages. Therefore, trophozoites (∼24 hpi) were exposed to the lead compounds at 10 x EC_50_ of growth for four hours before being washed out and allowed to proceed to the following cycle, where they were assessed for growth via Nluc activity. It was found that five of the invasion inhibitory compounds (MMV676877, MMV637229, MMV688279, MMV019721 and MMV020512) and one of the egress inhibitors (MMV016838), decreased trophozoite growth with invasion inhibitors MMV637229 and MMV688279 causing a significant reduction in growth when compared with control compound heparin (Figure 5B). This corroborated the results from the purified merozoite invasion assay which demonstrated that not all of these compounds specifically block merozoite invasion, thereby alluding to inhibitors that affect general processes in the parasite that may be required at the time of invasion. As MMV020291 and MMV006833 did not inhibit trophozoite growth and demonstrated invasion inhibitory activity in the purified merozoite invasion assay, we next investigated which stage of invasion was being blocked by these compounds.

### Live cell imaging of the two lead invasion inhibitors, MMV020291 and MMV006833, indicate MMV020291 blocks successful merozoite invasion of RBCs and MMV006833 slows down the invasion process, arresting ring formation

In order to gain an understanding of which stage of invasion might be affected by MMV020291 and MMV006833, we assessed the kinetics and physical morphology of *P. falciparum* invasion of RBCs by live cell microscopy using methods that had been previously developed (23) (Supplementary Videos 3-5). For consistency we used the same Hyp1-Nluc 3D7 parasite line as the invasion screen, with 0.1% DMSO as the vehicle control where we filmed and analysed 10, 11 and 12 egress events for MMV020291, MMV006833 and DMSO treatment, respectively. After each schizont rupture, in the presence of 10 µM MMV020291 or 2 µM MMV006833 (∼10 x EC_50_), the number of merozoites which adhered to neighbouring RBCs for ≥ 2 seconds per egress was comparable to DMSO (DMSO mean: 24.0, MMV02291 mean: 25.3, MMV006833 mean: 25.3) (Figure S5A). This indicated the compounds had not reduced the initial adhesiveness of the merozoites (note that merozoites may contact more than one RBC). Merozoites that maintain contact with their target RBCs then typically proceed to deform the surface of the RBC they attempt to invade. The time of initial merozoite contact to start of RBC deformation and duration of deformation upon MMV020291 treatment was comparable to the DMSO control (Figure 6A.i, S5B). In contrast, MMV006833 significantly delayed the merozoite’s ability to induce RBC deformation but did not affect the duration of deformation (Figure 6A.i, S5B).

**Figure 6.**
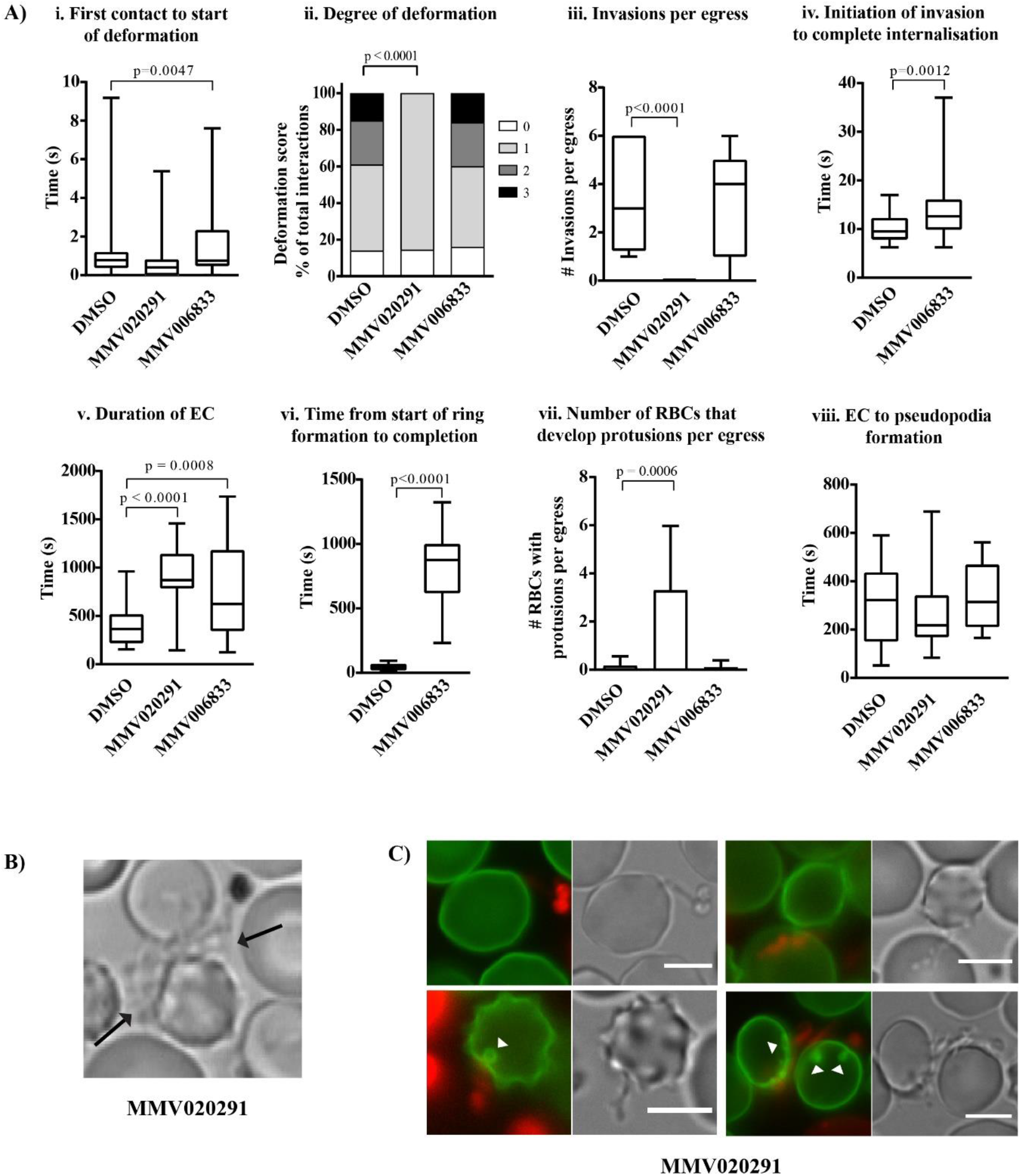
Live cell microscopy reveals MMV020291 blocks merozoite invasion and MMV006833 significantly slows down the invasion process and arrests conversion into ring stages. **A)** i. Analysis performed on live cell microscopy videos of parasites treated with 10 µM MMV020291 and 2 µM MMV006833 demonstrated an increase in the time taken for MMV006833 treated merozoites to cause deformation in the RBC membrane after first contact when compared with vehicle control, 0.1% DMSO. (ii) MMV020291 treated merozoites markedly lacked the ability to deform RBC membranes when compared with DMSO. (iii) MMV020291 revealed complete inhibition of merozoite invasion in contrast to DMSO and MMV006833 successful invasions. (iv) MMV006833 treated merozoites significantly increased the time taken from initialisation of the invasion event until complete internalisation within the RBC. (v) Treatment of merozoites with both invasion inhibitory lead compounds lead to significant prolonging in the duration of RBC echinocytosis. (vi) MMV006833 caused a significant increase in the time taken from the start of ring formation to completion. (vii) MMV020291 caused a significant number of protrusions to form at site of merozoite contact with target RBCs after egress events occurred. (viii) There were no differences observed between time taken from echinocytosis to pseudopodia formation for both DMSO and MMV006833 treatment after merozoite invasion and time taken from echinocytosis to external pseudopodia formation on the RBC surface with MMV020291 treatment. **B)** Frame of a live cell microscopy video showing a ruptured schizont treated with 10 µM MMV020291 and the merozoites that attempted but could not complete invasion forming protrusions extending from the point of contact with the RBC (black arrows) (Supplementary video 4). **C)** Brightfield and fluorescence images of failed merozoite invasions following 10 µM MMV020291 treatment. The RBCs were stained with BODIPY FL C12-Sphingomyelin (green) and the merozoites with BODIPY TR ceramide (red) which demonstrated the protrusions were derived from merozoite material. A green “punctum” was also sometimes observed on the RBC at site of merozoite contact (white arrow heads). 12, 10 and 11 schizont egress events were filmed and analysed for DMSO, MMV020291 and MMV006833 treatment, respectively. Images and videos analysed using ImageJ. Statistical analysis performed on GraphPad Prism using unpaired t tests (A.i, iii-viii) and Chi-square contingency (A.ii). No bar indicates not significant. EC= echinocytosis. Scale bars indicate 5 µm.

Next, the degree of RBC deformation caused by merozoite contact following compound treatments was scored to assess receptor-ligand interactions during early stages of *P. falciparum* invasion of RBCs where the intensity of RBC deformation is positively correlated with successful invasion (23). Merozoite contact with no RBC deformation was scored as ‘zero’ and strong deformation with the merozoite wrapping the RBC around itself was scored ‘three’ with intermediate effects scoring ‘one’ and ‘two’ (23). In the presence of MMV020291 there was a significant decrease in the deformation score whilst degree of deformation remained unchanged with MMV006833 treatment (Figure 6A.ii). A further qualitative observation was that treatment with MMV020291 appeared to reduce gliding motility of the merozoites across the RBC surface (Supplementary Video 3, DMSO and Supplementary Video 4, MMV020291).

Although MMV020291 treated merozoites appeared to attempt to invade their target RBCs, none completed the invasion process to achieve complete internalisation into the RBC. This is in contrast to an average of 3.4 and 3.3 invasions per schizont egress in the DMSO control and MMV006833 treatments, respectively (Figure 6A.iii). Although, MMV006833 treated merozoites invaded, they took significantly longer to penetrate their RBCs than the DMSO control (14.0 s vs. 10.1 s respectively, p=0.0012, Figure 6A.iv).

Merozoite invasions typically cause their target RBCs to rapidly undergo echinocytosis, a process where they develop a stellate appearance which returns to a normal biconcave shape after several minutes, by which time the merozoite has differentiated into a ring (23, 28). Although, MMV020291 treated merozoites did not successfully invade, they still triggered echinocytosis in an average of 4.0 RBCs per egress. This was not significantly different to the average 2.5 and 2.8 RBC echinocytosis events per egress in the DMSO control and MMV006833 treatment, respectively (Figure S5B, p =0.07 (DMSO and MMV020291)).

Even though MMV020291 and MMV006833 treatment still triggered RBC echinocytosis, the compounds greatly prolonged the echinocytosis period (903.6 and 795.8, respectively) compared to the DMSO control that saw an echinocytosis period of an average of 404.1 seconds (Figure 6A.v). Note that the echinocytosis periods for MMV compound treatments are an underestimate because the echinocytosed RBCs had often not recovered their normal shape by the end of the 20 minute filming period.

After invasion was complete, merozoites began to differentiate into amoeboid, ring stage parasites several minutes later. The process started with the growth of an arm-like projection or pseudopod from the internalised merozoite before full differentiation into an amoeba. In the DMSO control, ring conversion was completed in most invasions within one minute (Figure 6A.vi, Supplementary Video 3, black arrow), whereas ring formation appeared to be greatly slowed down or even arrested with MMV006833 treatment (mean: 803.9 seconds, Figure 6A.vi, p<0.0001, Supplementary Video 5, black arrows). The time taken for ring formation after MMV006833 treatment was an underestimate of the severity of the defect, since the 20 minute filming period frequently ended before ring formation was complete.

After echinocytosis had commenced following MMV020291 treatment, pseudopodial protrusions began to appear on the outside of RBCs where the merozoites had failed to invade (Figure 6B). Here, 100% of egress events and stalled invasions produced protrusions on at least one RBC, with a maximum of eight different RBCs developing protrusions after a single egress event (Figure 6A.vii). This is probably an underestimate of the pseudopod formation since many of the merozoite contacted RBCs had more than one protrusion. The formation of pseudopods from failed invasions was also occasionally observed with merozoites treated with DMSO and MMV006833, where 16.6% and 9.1% of egress events produced a single protrusion on the surface of a single RBC, respectively (Figure 6A.vii).

To gain an indication that the pseudopods formed from MMV020291 blocked invasions were equivalent to the formation of pseudopods during ring formation, we compared the duration of echinocytosis to pseudopodia formation for MMV020291 with DMSO and MMV006833 treatments. Here, no significant difference was observed (Figure 6A.viii, DMSO mean: 324.2 s, MMV020291 mean: 261.0 s, MMV006833 mean: 332.0 s), indicating that the protrusions formed after MMV020291 treatment might emanate from merozoites that had begun to differentiate into rings on the outside of the RBCs they had failed to invade.

In order to confirm that these pseudopods observed after MMV020291 treatment were parasite-derived, rather than originating from the RBC, purified schizonts and RBCs were stained with fluorescent green and red bodipy membrane dyes, respectively. This revealed that the MMV020291 induced pseudopods were red and therefore merozoite derived, either as a result of cell lysis or aberrant differentiation into ring-like parasites at the RBC surface following invasion failure. The fluorescent green RBC dye also revealed a distinct circular or “punctate structure” at the RBC invasion site, possibly originating from failed PV formation (Figure 6C, white arrows).

## Discussion

In this study, we have shown that by using the bioluminescent reporter protein, Nluc, it was possible to effectively screen compound libraries for inhibitors of parasite egress and invasion of RBCs in a microplate-based manner. Using this technique, we screened the MMV Pathogen Box and identified 15 compounds that inhibit parasite egress and 24 invasion-specific inhibitory compounds. We independently sourced 11 of these compounds and investigated their activity on *P. falciparum* using various growth and invasion assays, in addition to live cell microscopy.

After performing these validation assays, we grouped these compounds into one of five following categories; 1) blockers of late schizont maturation (MMV011765, MMV016838), inhibitor of the breakdown of the iRBC membranes and merozoite egress (MMV676881), direct blocker of merozoite invasion of RBCs (MMV020291), 4) inhibitor of the invasion process and ring development (MMV006833) and 5) general growth inhibitors that have a strong effect on invasion (MMV676877, MMV637299, MMV688279, MMV687794, MMV019721, MMV020512) (Figure 7).

**Figure 7.**
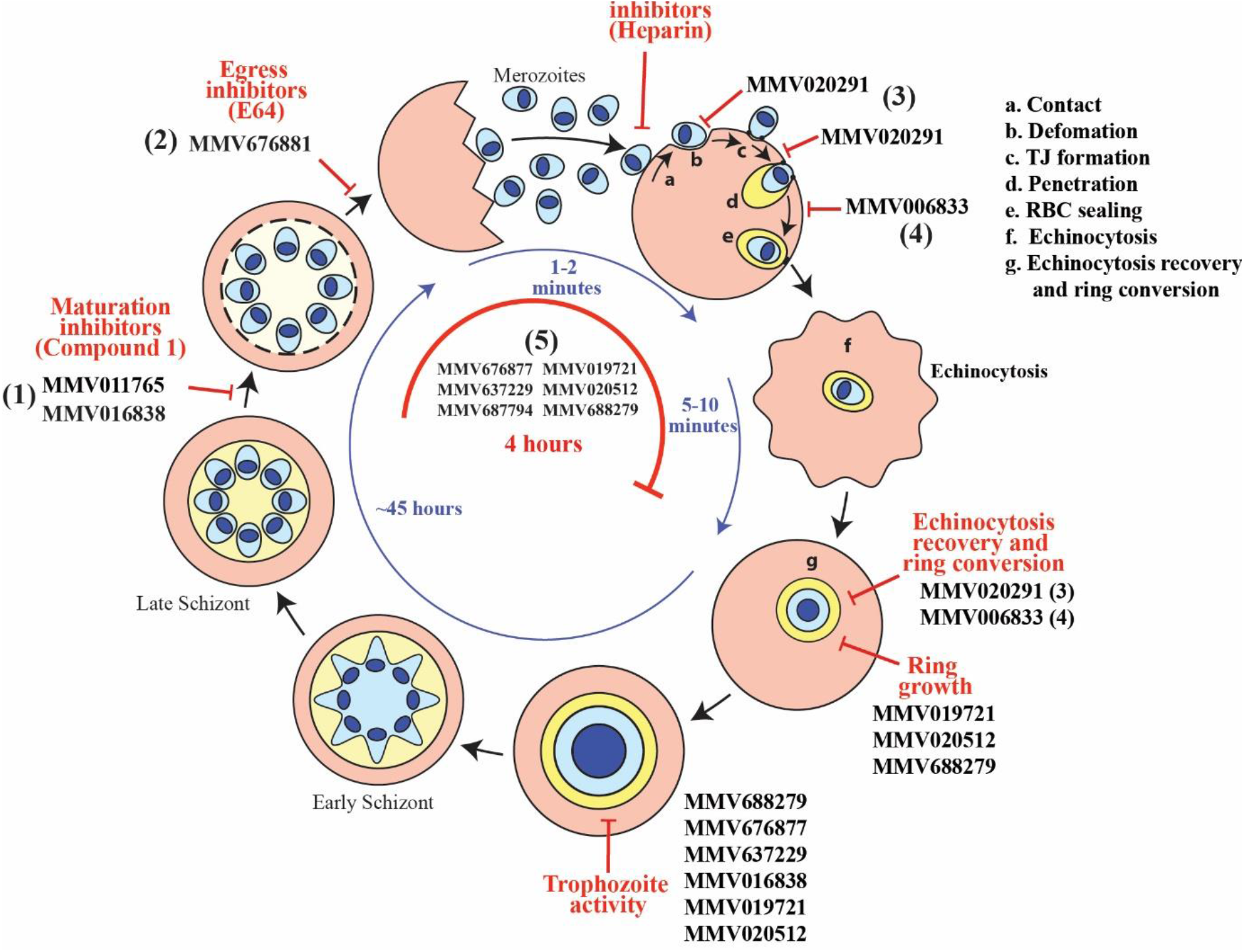
Summary of key effects of hit compounds that inhibit egress and invasion. **A)** Schematic of proposed locations where MMV Pathogen Box compounds act to inhibit egress and invasion. MMV011765 and MMV016838 act to inhibit schizont maturation, resembling treatment with the PKG inhibitor, Compound 1 (C1) (1). MMV676881 induces the formation of parasitophorous vacuole membrane enclosed merozoite structures that inhibits the release of merozoites, acting similarly to the protease inhibitor, E64 (2). MMV020291 blocks merozoite invasion by inhibiting penetration and prevents RBC recovery from echinocytosis (3) MMV006833 slows down the invasion process, prevent RBC echinocytosis recovery and blocks ring formation (4). MMV676877, MMV019721, MMV637229, MMV020512, MMV687794 and MMV688279 do not specifically block merozoite invasion or egress but may induce growth defects that indirectly hinders the invasion process (5).

The two Pathogen Box egress inhibitors had characteristics of E64 and C1, compounds that target cysteine proteases and PKG, respectively. PfPKG has been shown to be a master regulator of egress by sitting at the top of a cascade of events that culminate in merozoite release (11, 54, 55). Treatment of schizonts with MMV011765 and MMV016838 induced a developmental arrest similar to C1 treatment. However, MMV011765 appears to be irreversible suggesting that this compound permanently binds to its target or prevents a time sensitive event that cannot be resumed in order to produce viable merozoites.

The egress inhibitory compound, MMV676881, was originally identified by the Nluc screen as a putative invasion inhibitor since the Nluc enzyme was released from the schizonts, suggesting that schizont rupture had occurred. Giemsa stained smears, however, revealed the compound trapped merozoites inside unruptured iRBCs. We observed that the broad-spectrum cysteine protease inhibitor, E64, acts similarly to MMV676881 in the Nluc invasion assay by not preventing Nluc release. A probable explanation for the misclassification of MMV676881 is that the RBC membrane had become leaky to the Nluc reporter protein in mature schizonts, allowing it to escape into the growth media. This agrees with findings that demonstrate RBCs are permeable to small molecules, including Nluc reporter proteins, at late schizont stages in the presence E64 (54, 56, 57).

MMV676881 likely targets cysteine proteases as it was originally identified as an inhibitor of cruzain, a papain-like cysteine protease present in *Trypanosoma cruzi* (58, 59). Cysteine proteases have been shown to be crucial for schizont egress in *P. falciparum* through the use of E64 (60), as well as genetic manipulations of cysteine proteases such as serine rich antigen 6 (SERA6) whereby a conditional knockdown revealed it was essential for RBC membrane rupture of schizonts by reducing cytoskeletal stability (9, 60). Live cell microscopy and functional assays performed in this study indicated MMV676881 potentially acted as a *P. falciparum* cysteine protease inhibitor since the PEMS it produced could be mechanically broken to release invasion competent merozoites.

MMV020291 and MMV006833 were the most invasion specific inhibitors we identified in the screen. When schizonts were treated with MMV006833, MMV020291 or DMSO, no differences were observed in the number of merozoites which contacted RBCs per egress and their duration of deformation. This indicated that low affinity early interactions mediated by MSP1 and its associated proteins are likely to be unaffected by either compound (18). RBC deformation was, however, weaker with MMV020291 treatment and did not progress to strong deformation, defined by a merozoite pushing a furrow into the RBC surface. This indicates there may be a lack of the more intense and complex levels of deformation mediated by EBA and PfRhs interactions (61, 62). These observations are consistent with MMV020291 inhibiting the discharge of micronemes and/or rhoptries (63, 64).

In addition to the lack of extreme deformation, merozoites treated with MMV020291 did not appreciably migrate across the surface of the RBC, a behaviour known as gliding motility that has been observed in normal merozoite-RBC interactions (23, 65). The actin-myosin motor is critical for deformation, motility and RBC penetration whereby merozoites treated with actin polymerisation inhibitor, cytochalasin D, have prevented migration over RBCs and failure of invasion (23). MMV020291 could therefore be inhibiting the actin-myosin motor, although some functionality must be retained since many merozoites were seen to push into the RBC at the point at which echinocytosis began within the timeframe of normal invasion and were also observed to partially form a PV.

Although MMV020291 appears to inhibit strong RBC deformation, a function mediated by EBAs and PfRhs, it does not appear to inhibit the downstream-acting PfRh5. This protein is the only non-redundant PfRh and binds to the RBC receptor, basigin, that probably activates secretion of the RON complex to enable AMA1-RON2 tight junction formation (23, 66, 67). Antibodies to PfRh5, or basigin, inhibit merozoite invasion but pre-invasion, deformation and reorientation processes remain unaffected (23, 29). This is not what we observed following MMV020291 treatment suggesting PfRh5 functions are not the target. PfRh5 inhibition also prevents echinocytosis from occurring suggesting MMV020291 also functions downstream of this protein since echinocytosis clearly occurs following MMV020291 treatment. In this respect, MMV020291 more closely mimics the effects of blocking the AMA1 and RON complex interaction, where merozoites remain attached after echinocytosis was initiated and RBC recovery to its normal biconcave shape was greatly delayed (29, 68). MMV020291 therefore inhibits a range of early and late acting invasion functions making it unclear what the possible target may be.

We observed MMV020291 treated merozoites had formed protrusions at the site of contact with the RBC surface. These protrusions were stained the same fluorescent dye that was used to label the merozoites, indicating that they originated from parasite material. These protrusions may therefore be merozoites that have differentiated into ring-stage parasites on the outside of the RBC as newly invaded merozoites often produce mobile, pseudopodial extensions within the iRBC prior to ring differentiation which can be observed in a normal invasion event in Supplementary Video 3. Live cell microscopy of failed invasion events have not been previously described to form protrusions as recently observed with PKA and adenylate cyclase beta gene disruptions (69). Further support that MMV020291 treatment causes unsuccessful merozoites to differentiate into rings is that there was no significant difference observed between the time taken for pseudopod formation to become visible on the surface of MMV020291 treated RBCs compared to normally invaded RBCs. The small, partially formed PV-like structure that was visible in some of the failed invasion sites of MMV020291 treated merozoites indicates parasite induced modification of the RBC surface had occurred. These PV-like structures could be the equivalent of the whorl-like membranous structures that has been seen to form at the RBC surface from rhoptry contents when merozoite invasion was arrested with cytochalasin D (70).

In contrast to MMV020291, MMV006833 treatment did not affect the merozoite’s ability to deform RBCs, form tight junctions and invade RBCs, indicating that the release of the rhoptries, micronemes and associated interactions with RBC receptors were unaffected. Whilst MMV006833 treated merozoites were able fully enter their RBCs, the invasion process was significantly slower than normal invasion events suggesting that the actin-myosin motor may be affected. However, treatment with MMV006833 did not affect the merozoite’s ability to cause severe deformation, which indicates that the actin-myosin motor can still function as treatment with cytochalasin D has previously been shown to block deformation (23). Once invaded, the merozoites treated with MMV006833 were observed to arrest in at the pseudopod stage, unable to differentiate into ring-stage parasites and this may be the mechanism by which this compound blocks growth. There is little known about the mechanisms underlying merozoite differentiation into ring stage parasites post invasion and identifying the target of MMV006833 may further the study of this process.

The remaining six compounds (MMV676877, MMV637229, MMV688279, MMV687794, MMV019721 and MMV020512) from the lead list we have termed “general growth inhibitors” as whilst they inhibited schizont progression to ring-stage parasites in the primary Nluc screen, they did not specifically block parasite invasion. This was corroborated by growth assays that determined five out of six of these compounds have inhibitory activity across other stages of the lifecycle (MMV676877, MMV637229, MMV688279, MMV019721 and MMV020512). Supporting this, a screen of the MMV Pathogen Box to identify stages of compound activity in the *P. falciparum* RBC 48-hour lifecycle by using DNA content as a marker of stage arrest, classified these five compounds as either arresting parasite growth without DNA replication (i.e. ring-stage) or halting growth at trophozoite stage without sufficient DNA replication (71). It is therefore likely, some of these compounds could disrupt general cellular pathways such as DNA regulation or transcription and translation machinery. It has been demonstrated in schizonts that 10% of the parasite genome has a high level of transcription, with genes encoding for proteins such as MSPs and rhoptries being upregulated in late schizogony (72, 73). Four hit compounds from the primary Nluc screen targeting elongation factor 2 (EF2) and DNA binding agents were removed for possessing ring stage activity and lacking novel targets in the compound triaging process (43, 46, 49). This demonstrates that agents that block DNA regulation may inadvertently inhibit essential invasion genes from being transcribed, thereby inducing invasion inhibitory phenotypes. Whilst an *in silico* analysis of potential molecular docking candidates to PfCDPK5 identified MMV020512 as a hit compound (74), we have failed to observe any inhibitory phenotypes that have previously been seen with PfCDPK5 knockdowns, namely the development of PEMS (10, 57).

The sixth of these general growth inhibitors, MMV687794, was identified as an inhibitor that arrested late trophozoite/schizont after DNA replication occurred in the aforementioned study (71) and whilst, it did not directly inhibit invasion in the purified merozoite assays, we saw no growth defects at other stages. This could indicate that this compound may block targets upstream of invasion that are required to “prime” the merozoite for invasion in the schizont stage such as proteins that undergo proteolytic processing in schizonts like AMA1, MSP1 and the rhoptry associated protein, RAP1 (75–80).

In conclusion, we have identified three specific merozoite egress, one RBC invasion inhibitor and one inhibitor that slows invasion and arrests ring development, in addition to several other general growth inhibitors that strongly act during the invasion stages. These inhibitors, along with their novel mechanisms of action, could complement current antimalarials which generally act on intracellular parasites during their growth phase. It will be important to identify the target proteins of the egress and invasion inhibitor compounds because this will inform structure-activity relationship based drug design to improve the compounds’ potencies. Once their targets are known, these compounds could also act as useful tools to further dissect molecular details of egress and invasion processes in the parasite.

## Methods

### Parasite culture and strains

*Plasmodium falciparum* parasites were continuously cultured as previously described (81) in human RBCs (Australian Red Cross Blood Bank, Type O) at 4% haematocrit in supplemented RPMI media (RPMI-HEPES, 0.2% NaHC0_3_, 5% heat-inactivated human serum [Australian Red Cross], 0.25% AlbumaxII [GIBCO], 0.37 mM hypoxanthine, 31.25 µg/mL Gentamicin) at 37°C. An exported Nluc parasite line was used as previously described (33), which was generated by transfecting a Hyp1-Nluc plasmid under the control of an *ef1α* promoter into 3D7 *P. falciparum* parasites (82). Uptake of the plasmid was selected for and maintained by 2.5 nM WR99210. This parasite line was used for all experiments in the study.

### Compounds

Pathogen Box compounds were obtained from the Medicines for Malaria Venture (MMV) and consisted of 400 compounds at 10 mM dissolved in dimethyl sulfoxide (DMSO) (https://www.pathogenbox.org/). Compounds were diluted to 100 µM and aliquoted into 96 well plates at −80°C for long-term storage. Further quantities of MMV011765, MMV637229, MMV020291, MMV006833, MMV688279, MMV676877 and MMV676881 were provided by MMV and MMV016838 (MolPort-001-614-591), MMV020512 (MolPort-004-158-754), MMV019721 (MolPort-004-102-322) and MMV687794 (MolPort-002-553-011) were purchased from Molport. R1 peptide (a kind gift from Alan Cowman) and porcine heparin (Sigma) were dissolved in RPMI media. Chloroquine (Sigma) and E64 (Sigma) were dissolved in water. Artemisinin (Sigma) and Compound 1 (custom made) and all other compounds were dissolved in DMSO.

### Nanoluciferase invasion assay and statistical validation for HTS

Parasites were grown to late stage schizonts and isolated using a Percoll density gradient whereby culture was added to 60% buffered Percoll solution in supplemented RPMI (10 mM NaH_2_PO_4,_ 143 mM NaCl, Percoll [GE Healthcare Bio-Sciences]). Purified schizonts at 1-2% parasitemia were added to RBCs with a final haematocrit of 1%. Drugs were administered at concentrations of 0.02%, 4 µM and 100 µg/mL for DMSO, C1 and heparin, respectively. Plates were incubated at 37°C for 4 h and the growth media was removed to measure Nluc released upon schizont egress (see below). Cells were treated with sorbitol and three washes were performed before incubating the culture at 37°C for a further 24 h until parasites were >24 hpi. Whole cells were lysed to measure intracellular Nluc, corresponding with initial invasion rate (see below). To account for contamination of ring stage parasites that failed to be removed from Percoll gradient and schizonts that failed to be removed by sorbitol treatment, a background control was included whereby Percoll purified cultures were kept at 4°C during the first 4 h incubation and subsequently treated with sorbitol before further incubation at 37°C for 24 h.

### Measuring Nanoluciferase activity

Whole cells (5 µL) at 1% hematocrit (invasion) or growth media (egress) was dispensed into white 96 well luminometer plates and 45 µL of 1 x NanoGlo Lysis Buffer containing 1:1000 NanoGlo substrate (Promega) was injected into wells. Relative light units (RLU) was measured by a CLARIOstar luminometer (BMG Labtech).

### Analysis

The percent of invasion/egress was determined by subtracting the RLU of background control and normalising all values to the average RLU of the untreated control. The mean and standard deviation of 60 total wells (15 wells per plate over four biological replicates) for each drug treatment was used to calculate Z’ scores using the following equation as per (36):

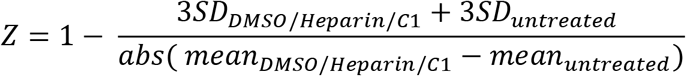

### Screening the Pathogen Box

The Nanoluciferase invasion assay was used as described above. Pathogen Box compounds were diluted to a final concentration of 2 µM with concentrations of DMSO, C1 and heparin as listed above. Chloroquine was included at 75 nM. Each plate contained 40 compounds in duplicate with control compounds. Percentage of invasion and egress in the presence of compounds was normalised relative to the mean of DMSO (100% egress or invasion rate) that was averaged across three biological replicates.

### Counter screen

Parasites were grown to 24 hpi and adjusted to a final haematocrit of 1% with 1-2% parasitemia. Cells were lysed in 1 X Nanoglo Lysis Buffer (Promega) and lysate was added to 10 µM of the Pathogen Box compounds and incubated for 10 minutes at 37°C. Nanoluciferase activity was measured as above with 45 µL of lysate dispensed into 96 well white luminometer plates and 5 µL 1:100 NanoGlo substrate injected into wells. Nluc activity in presence of compounds was normalised relative to 0.1% DMSO (100% Nluc activity).

### Early ring inhibition assay

This was performed as per the Nluc invasion assay with Pathogen Box compounds tested at a concentration of 2 µM. Chloroquine and artemisinin were included at concentrations of 75 nM and 25 nM, respectively. Duplicate wells for each compound were set up, one that received drug treatment during the egress/invasion window and one that remained untreated.

After the 4 h incubation, following sorbitol treatment and washes, the well that was untreated received drug treatment for a further 4 h before being washed 3x. At 24 hpi cultures were lysed and Nluc activity measured as described above. Percentage of invasion and early ring growth in the presence of compounds was normalised relative to 0.02% DMSO (100% invasion or early ring growth).

### Egress inhibition reversibility assay

Schizonts were Percoll purified as described above and cultured in 1% haematocrit of 1-2% parasitemia. To this, C1, MMV011765, MMV016838 and DMSO were added at final concentrations of 4 µM, 10 µM, 10 µM and 0.1%, respectively. Plates were incubated for 2 h at 37°C before growth media was collected. Pellets were washed 3x and incubated for a further 4 h at 37°C before growth media was collected. Nluc released in the growth media was measured as described above. Egress was normalised to relative to 0.1% DMSO.

### Merozoite invasion assay

Adapted from (51–53) where parasites were grown to schizonts, magnet purified and treated with 10 µM E64 (or 10µM MMV676881) for 4-6 hours until >50% had become PEMS. Schizonts were washed with supplemented RPMI before being mechanically disrupted by passage through a 1.2 µm filter and distributed into a 96-well plate with a final hematocrit of 1% and ∼10 x EC_50_ of growth of each drug (10 µM MMV020291, 3 µM MMV006833, 3 µM MMV668279, 7 µM MMV676877, 3 µM MMV687794, 5 µM MMV020512, 5 µM MMV019721, 10 µM MMV637229 (5 x EC_50_)). The 96-well plate was shaken at 400 rpm for 30 minutes at 37°C. Pellets were washed 3x and put back into culture for 24 h before Nluc activity was measured as described above. Invasion was normalised relative to 0.1% DMSO.

### Trophozoite growth assays

Hyp1-Nluc parasites at ∼24 hpi were adjusted to 1% parasitemia and 1% haematocrit in a 96-well plate before being exposed to 10 x EC_50_ of Pathogen Box compounds for 4 hrs. Plates were then washed 3x in supplemented RPMI before being put back into culture until the following cycle. Growth was calculated at ∼24 hpi by measuring Nluc activity as described above. C1, heparin, chloroquine and artemisinin were included as controls at 4 µM, 100 µg/mL, 75 nM and 25 nM, respectively. Values were normalised to 0.1% DMSO as a vehicle control.

### Determination of EC_50_s for parasite invasion and egress of RBCs

Compounds were serially diluted from 10 µM and rates of parasite egress and invasion were measured by the Nluc invasion assay as described above. Percentage of invasion and egress in the presence of compounds was normalised to 0.1% DMSO (100% invasion or egress rate). EC_50_ curves were generated from GraphPad Prism 7 using a nonlinear regression.

### Live cell microscopy

Live cell microscopy was performed as described in (23, 83) on a Zeiss Cell Observer widefield fluorescent microscope equipped with a humidified gas chamber (1% O_2_, 5% CO_2_ and 94% N_2_) at 37°C. Eight well LabTek chambered slides were filled with 200 µL of parasite culture, diluted to a final 0.1% haematocrit in RPMI media and treated with 10 x EC_50_ of compound. Mature schizonts that appeared to rupture were imaged at two to four frames per second using the AxioCam 702 Mono camera for 20 minutes. Image and data analysis of cell behaviour was performed using ImageJ and GraphPad Prism with statistical tests between treated and vehicle control including unpaired t tests (for normally distributed data) and Man-Whitney t test (for non-normally distributed data).

### Bodipy staining

RBCs at 1% haematocrit were stained with bodipy sphingomyelin green (Life Technologies) diluted 1:500 in RPMI by incubating overnight at 37°C. Trophozoites were incubated with 1:1000 bodipy ceramide Texas red (Life Technologies) overnight in RPMI as described above but without human serum and decreased Albumax (0.125%). Approximately 12 h later, schizonts were Percoll purified as described above and stained RBCs were washed 2x in RPMI. Schizonts and RBCs were combined and then imaged as described above for live cell microscopy conditions.

## Acknowledgements

This work was supported by the Victorian Operational Infrastructure Support Program received by the Burnet Institute. We acknowledge Medicines for Malaria Venture (MMV) for providing access to the MMV Pathogen Box and the Australian Red Cross Blood Bank for the provision of human blood. M.G.D is a recipient of an Australian Government Research Training Program Scholarship, G.E.W a Peter Doherty - Australian Biomedical Fellowship, B.E.S a Development Grant 1113712, D.W.W and B.E.S NHMRC Project Grant APP1143974, and T.F.dK-W a NHMRC Senior Research Fellowship. B.E.S. is a Corin Centenary Fellow. D.W.W is a University of Adelaide Beacon Fellow. We thank Alan Cowman for providing the R1 peptide and Monash Micro Imaging for assistance with microscopy.

## Supplementary Figures

**Supplementary Figure 1.**
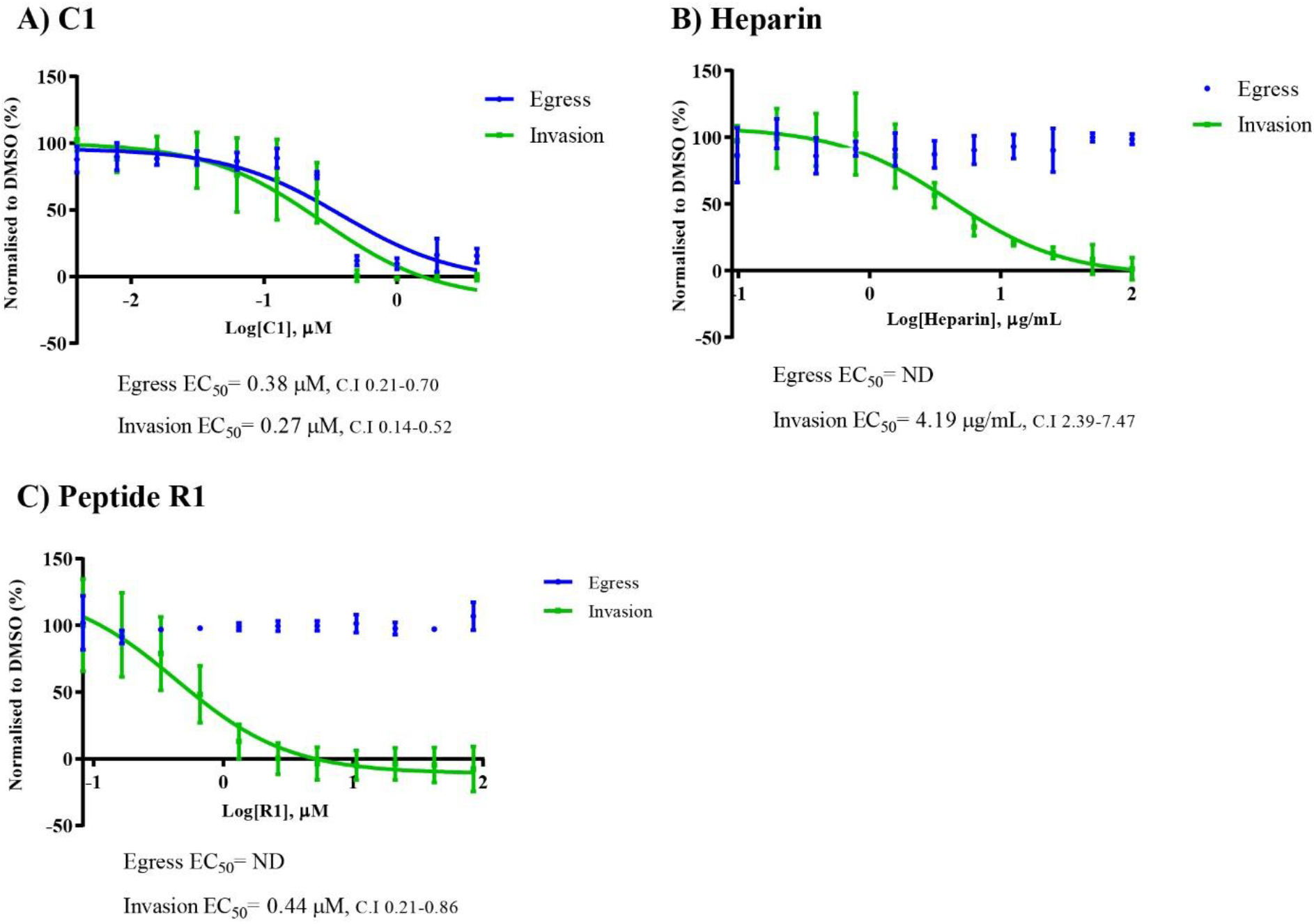
Egress and Invasion inhibitory compounds produce dose response curves for egress and invasion in the Nluc invasion assay. Graphs depict dose-response curves for the Nluc invasion assay of the following compounds: A) C1, B) Heparin and C) Peptide R1. Error bars represent standard deviation of 3 biological replicates whereby C.I indicates 95% confidence intervals for EC_50_ values which were derived from GraphPad Prism. ND= not determined.

**Supplementary Figure 2.**
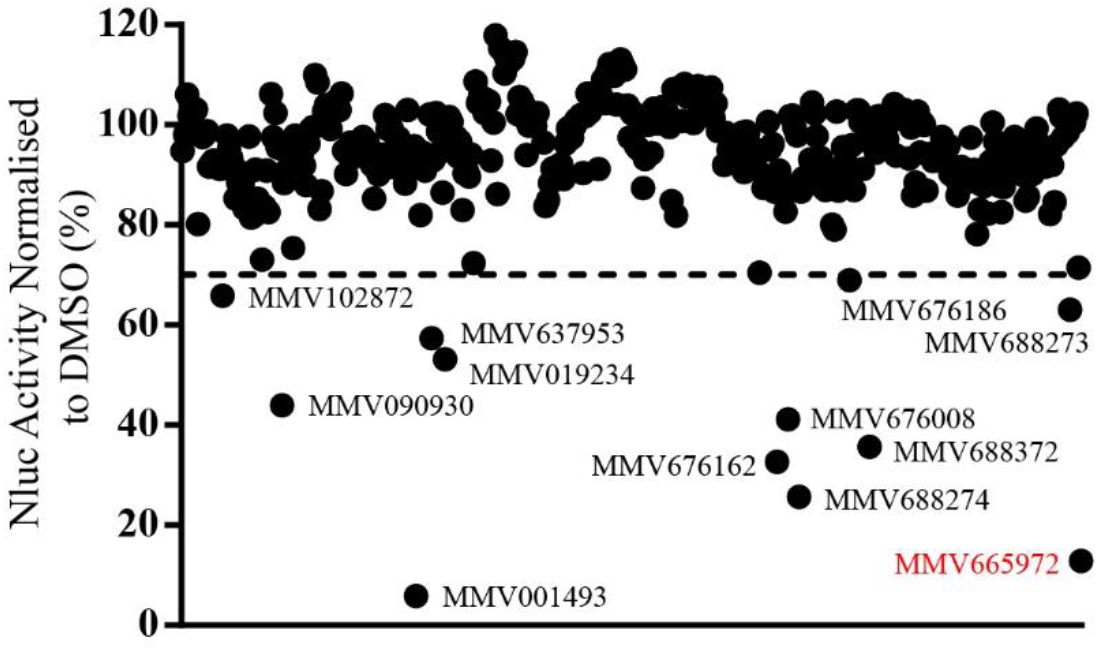
Counter screen of MMV Pathogen Box compounds. Nanoluciferase (Nluc) expressing parasites were lysed in a Triton-X buffer and exposed to 10 µM Pathogen Box compounds. Nluc activity was measured and eleven compounds were found to inhibit Nluc activity to <70%. MMV665972 was used as a control for Nluc inhibition as it has previously been found to inhibit Nluc activity (unpublished data). Each dot represents the mean of 2 biological replicates.

**Supplementary Figure 3.**
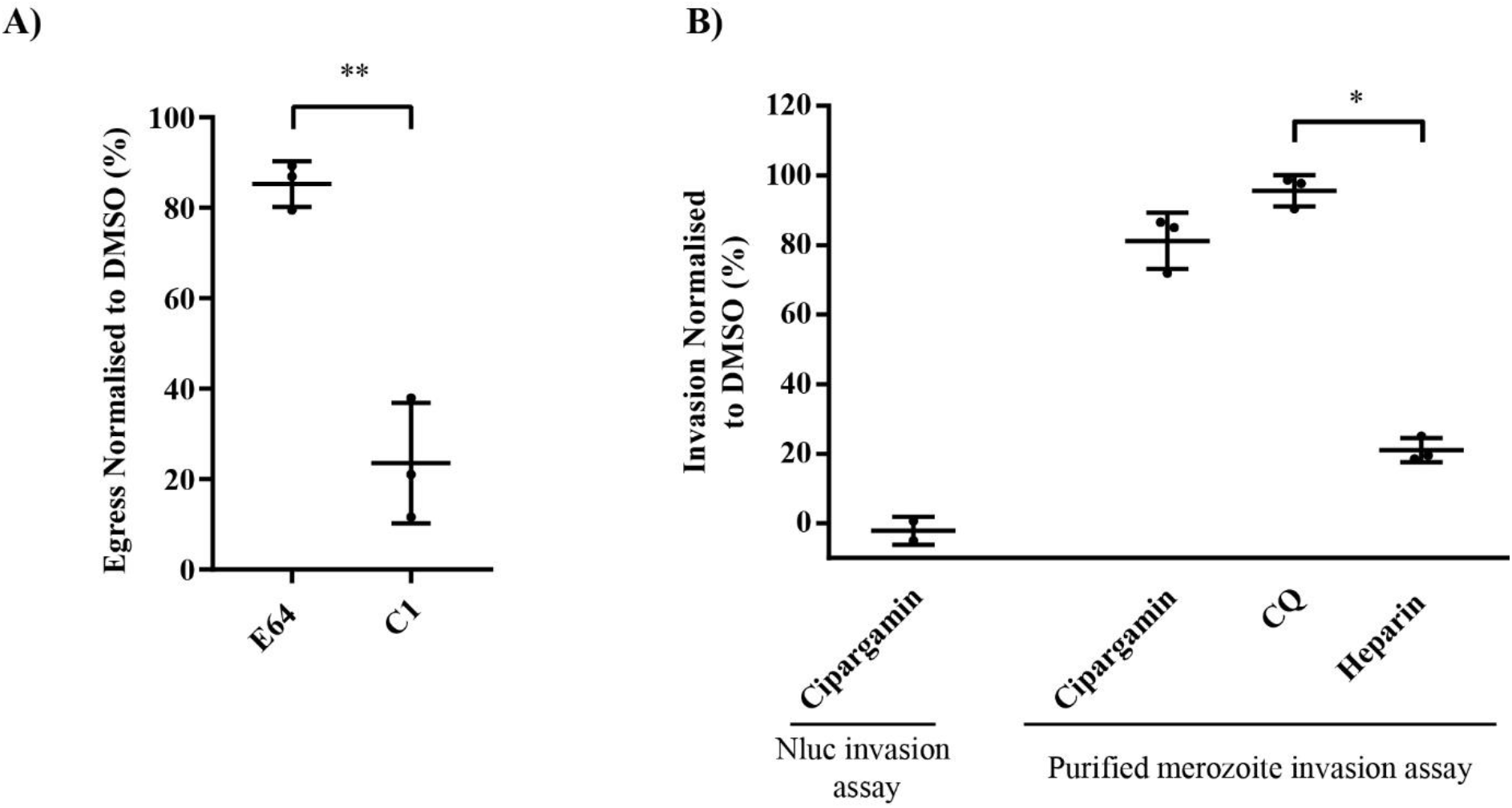
Egress inhibitor, E64, does not prevent release of Nluc from schizonts and PfATP4 inhibitor, cipargamin, does not inhibit invasion in purified merozoite assay. **A)** Using the Nluc invasion assay method, schizonts were treated with 10 µM of the egress inhibitor, E64, which did not inhibit the release of Nluc into the growth media when compared with 4 µM of the egress inhibitor, C1. B) The Nanoluciferase (Nluc) invasion assay demonstrated that PfATP4 inhibitor, cipargamin, inhibits invasion over the 4 hour window of schizont egress to merozoite invasion, however purified merozoite invasion assays showed that it did not prevent specific invasion into RBCs, in contrast to invasion inhibitor, heparin. Cipargamin, chloroquine (CQ) and heparin were used at concentrations of 20 nM, 75 nM and 100 µg/mL. Values have been normalised to 0.1% DMSO. Error bars represent the standard deviation of 3 biological replicates (A) and 2 and 3 biological replicates for Nluc invasion assay and purified merozoite invasion assay, respectively (B). Statistical analysis performed via unpaired t test (A) and one-way ANOVA in GraphPad Prism between CQ and heparin or cipargamin (B). * indicates p <0.05 and ** indicates p< 0.01. No bar indicates not significant.

**Supplementary Figure 4.**
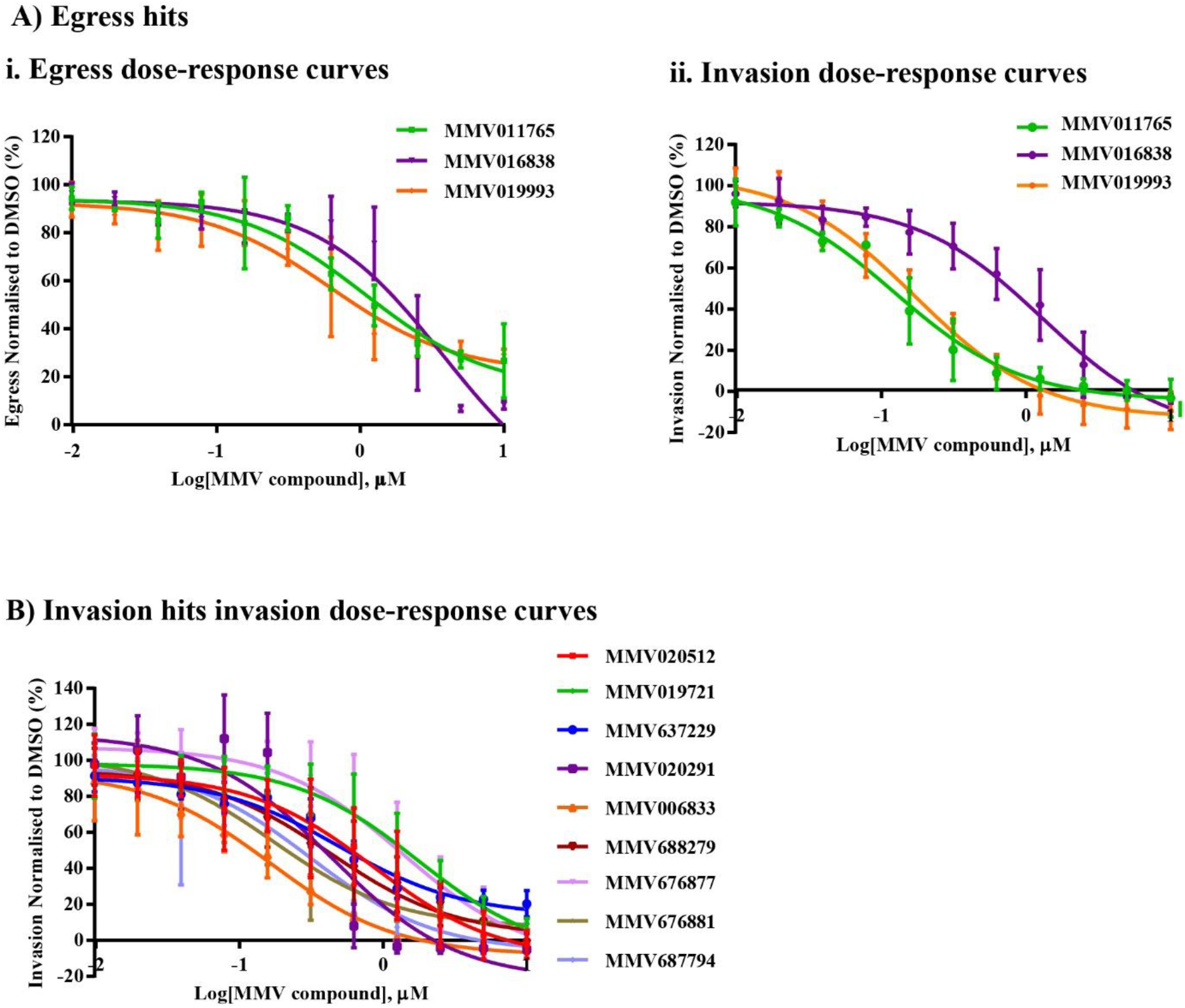
Dose-response curves for egress and invasion of lead compounds identified in the Nluc invasion screen. Lead compounds were tested at multiple concentrations in the Nluc assay in order to produce half maximal effective concentrations (EC_50_) for egress and invasion for compounds that were egress hits **(Ai, ii)** and invasion hits **(B)**. Error bars represent the standard deviation of 3 biological replicates whereby C.I indicates 95% confidence intervals for EC_50_ values. EC_50_ curves were generated from GraphPad Prism.

**Supplementary Figure 5.**
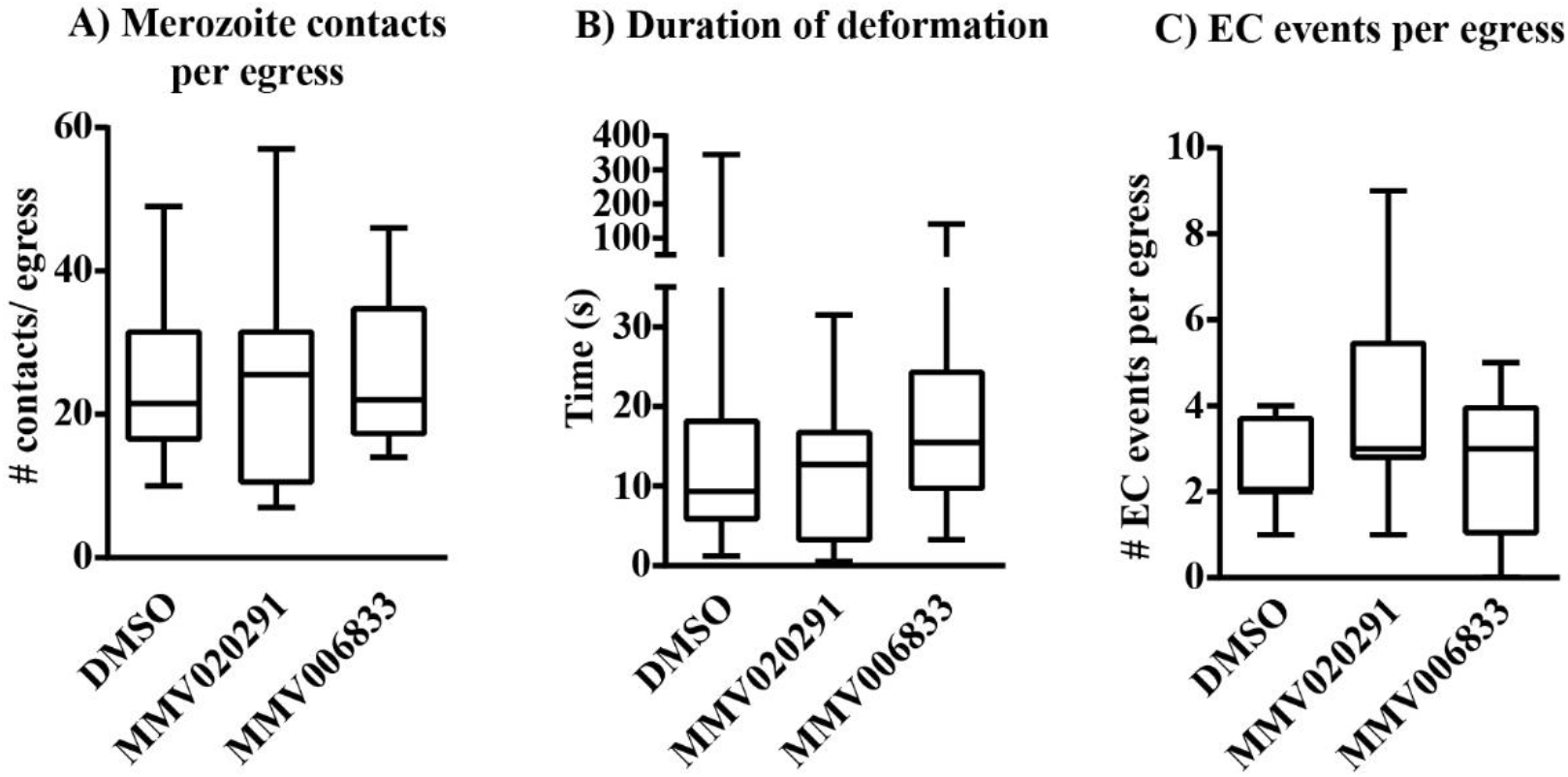
Live cell microscopy demonstrates lead invasion inhibitors, MMV020291 and MMV006833, do not affect merozoite contact, duration of deformation and echinocytosis after egress. Live cell microscopy movies of egress events were analysed for merozoite contacts per egress (A), duration of deformation (B) and echinocytosis events per egress (C) and there were no differences observed between the DMSO vehicle control and invasion inhibitory compounds, MMV020291 and MMV006833. 12, 10 and 11 schizont egress events were filmed and analysed for DMSO, MMV020291 and MMV006833 treatment, respectively. Images and videos analysed using ImageJ. Statistical analysis performed on GraphPad Prism using unpaired t tests. No bar indicates not significant. EC= echinocytosis.

**Supplementary Table 1. MMV Pathogen Box screening results for egress and invasion against *P. falciparum at 2 μM***

The 400 compound Pathogen Box invasion and egress values normalised to 0.02% DMSO vehicle control. SD indicates standard deviation of 3 biological replicates. Compounds highlighted in orange were hits identified as egress inhibitors by reducing egress rates <60% of DMSO control. Compounds highlighted in blue were hits identified as invasion inhibitors by reducing invasion rates <10% of DMSO control. Compounds highlighted in green were both hits of egress and invasion.

**Supplementary Video 1. 10 µM E64 treated schizonts produce PVM-enclosed merozoite structures that prevent merozoite egress**

**Supplementary Video 2. 4 µM MMV676881 treated schizonts produce a similar effect to E64 treatment that induce PVM-enclosed merozoite structures that inhibit merozoite egress**

**Supplementary Video 3. 0.1% DMSO treated schizonts egress and invade normally**

White arrows indicate successful invasions and the black arrow shows ring formation of invaded merozoite.

**Supplementary Video 4. 10 µM MMV020291 treated schizonts egress normally but inhibit the penetration of merozoites into RBCs and echinocytosis recovery**

Black arrows indicate merozoites that are attached to RBCs and unravel to form pseudopods.

**Supplementary Video 5. 2 µM MMV006833 treated schizonts egress normally but slow down the invasion process and arrest early ring formation**

White arrows indicate successful invasions and the black arrows show invaded merozoites that do not form rings.

## References

1. World Health Organization. 2018. World Malaria Report 2018.

2. Vijaykadga S, Rojanawatsirivej C, Cholpol S, Phoungmanee D, Nakavej A, Wongsrichanalai C. 2006. In vivo sensitivity monitoring of mefloquine monotherapy and artesunate– mefloquine combinations for the treatment of uncomplicated falciparum malaria in Thailand in 2003. Tropical Medicine & International Health 11:211–219.

3. Dondorp AM, Nosten F, Yi P, Das D, Phyo AP, Tarning J, Lwin KM, Ariey F, Hanpithakpong W, Lee SJ, Ringwald P, Silamut K, Imwong M, Chotivanich K, Lim P, Herdman T, An SS, Yeung S, Singhasivanon P, Day NPJ, Lindegardh N, Socheat D, White NJ. 2009. Artemisinin Resistance in *Plasmodium falciparum* Malaria. The New England journal of medicine 361:455–467.

4. Fairhurst RM, Dondorp AM. 2016. Artemisinin-resistant *Plasmodium falciparum* malaria. Microbiology spectrum 4:10.1128/microbiolspec.EI10-0013-2016.

5. Wickham ME, Culvenor JG, Cowman AF. 2003. Selective Inhibition of a Two-step Egress of Malaria Parasites from the Host Erythrocyte. Journal of Biological Chemistry 278:37658–37663.

6. Chandramohanadas R, Park Y, Lui L, Li A, Quinn D, Liew K, Diez-Silva M, Sung Y, Dao M, Lim CT, Preiser PR, Suresh S. 2011. Biophysics of Malarial Parasite Exit from Infected Erythrocytes. PLOS ONE 6:e20869.

7. Blackman MJ. 2008. Malarial proteases and host cell egress: an ‘emerging’ cascade. Cellular Microbiology 10:1925–1934.

8. Wirth CC, Pradel G. 2012. Molecular mechanisms of host cell egress by malaria parasites. International Journal of Medical Microbiology 302:172–178.

9. Thomas JA, Tan MSY, Bisson C, Borg A, Umrekar TR, Hackett F, Hale VL, Vizcay-Barrena G, Fleck RA, Snijders AP, Saibil HR, Blackman MJ. 2018. A protease cascade regulates release of the human malaria parasite *Plasmodium falciparum* from host red blood cells. Nature Microbiology 3:447–455.

10. Dvorin JD, Martyn DC, Patel SD, Grimley JS, Collins CR, Hopp CS, Bright AT, Westenberger S, Winzeler E, Blackman MJ, Baker DA, Wandless TJ, Duraisingh MT. 2010. A Plant-Like Kinase in *Plasmodium falciparum* Regulates Parasite Egress From Erythrocytes. Science (New York, NY) 328:910–912.

11. Hopp CS, Bowyer PW, Baker DA. 2012. The role of cGMP signalling in regulating life cycle progression of Plasmodium. Microbes and Infection / Institut Pasteur 14:831–837.

12. Sam-Yellowe TY. 1996. Rhoptry organelles of the apicomplexa: Their role in host cell invasion and intracellular survival. Parasitology Today 12:308–316.

13. Bannister LH, Hopkins JM, Fowler RE, Krishna S, Mitchell GH. 2000. A Brief Illustrated Guide to the Ultrastructure of *Plasmodium falciparum* Asexual Blood Stages. Parasitology Today 16:427–433.

14. Farrow RE, Green J, Katsimitsoulia Z, Taylor WR, Holder AA, Molloy JE. 2011. The mechanism of erythrocyte invasion by the malarial parasite, *Plasmodium falciparum*. Seminars in Cell & Developmental Biology 22:953–960.

15. Holder AA, Freeman RR. 1984. The three major antigens on the surface of *Plasmodium falciparum* merozoites are derived from a single high molecular weight precursor. The Journal of Experimental Medicine 160:624.

16. Lin CS, Uboldi AD, Marapana D, Czabotar PE, Epp C, Bujard H, Taylor NL, Perugini MA, Hodder AN, Cowman AF. 2014. The Merozoite Surface Protein 1 Complex Is a Platform for Binding to Human Erythrocytes by *Plasmodium falciparum*. The Journal of Biological Chemistry 289:25655–25669.

17. Vogt Anna M, Winter G, Wahlgren M, Spillmann D. 2004. Heparan sulphate identified on human erythrocytes: a *Plasmodium falciparum* receptor. Biochemical Journal 381:593–597.

18. Boyle MJ, Skidmore M, Dickerman B, Cooper L, Devlin A, Yates E, Horrocks P, Freeman C, Chai W, Beeson JG. 2017. Identification of Heparin Modifications and Polysaccharide Inhibitors of *Plasmodium falciparum* Merozoite Invasion That Have Potential for Novel Drug Development. Antimicrobial Agents and Chemotherapy 61.

19. Camus D, Hadley TJ. 1985. A *plasmodium falciparum* antigen that binds to host erythrocytes and merozoites. Science 230:553+.

20. Mayer DCG, Kaneko O, Hudson-Taylor DE, Reid ME, Miller LH. 2001. Characterization of a *Plasmodium falciparum* erythrocyte-binding protein paralogous to EBA-175. Proceedings of the National Academy of Sciences of the United States of America 98:5222–5227.

21. Lanzillotti R, Coetzer TL. 2006. The 10 kDa domain of human erythrocyte protein 4.1 binds the *Plasmodium falciparum* EBA-181 protein. Malaria Journal 5:100–100.

22. Li X, Marinkovic M, Russo C, McKnight CJ, Coetzer TL, Chishti AH. 2012. Identification of a specific region of *Plasmodium falciparum* EBL-1 that binds to host receptor glycophorin B and inhibits merozoite invasion in human red blood cells. Molecular and Biochemical Parasitology 183:23–31.

23. Weiss GE, Gilson PR, Taechalertpaisarn T, Tham W-H, de Jong NWM, Harvey KL, Fowkes FJI, Barlow PN, Rayner JC, Wright GJ, Cowman AF, Crabb BS. 2015. Revealing the Sequence and Resulting Cellular Morphology of Receptor-Ligand Interactions during *Plasmodium falciparum* Invasion of Erythrocytes. PLoS Pathogens 11:e1004670.

24. Cao J, Kaneko O, Thongkukiatkul A, Tachibana M, Otsuki H, Gao Q, Tsuboi T, Torii M. 2009. Rhoptry neck protein RON2 forms a complex with microneme protein AMA1 in *Plasmodium falciparum* merozoites. Parasitology International 58:29–35.

25. Lamarque M, Besteiro S, Papoin J, Roques M, Vulliez-Le Normand B, Morlon-Guyot J, Dubremetz J-F, Fauquenoy S, Tomavo S, Faber BW, Kocken CH, Thomas AW, Boulanger MJ, Bentley GA, Lebrun M. 2011. The RON2-AMA1 Interaction is a Critical Step in Moving Junction-Dependent Invasion by Apicomplexan Parasites. PLOS Pathogens 7:e1001276.

26. Dobrowolski JM, Sibley LD. 1996. Toxoplasma Invasion of Mammalian Cells Is Powered by the Actin Cytoskeleton of the Parasite. Cell 84:933–939.

27. Gonzalez V, Combe A, David V, Malmquist NA, Delorme V, Leroy C, Blazquez S, Ménard R, Tardieux I. 2009. Host Cell Entry by Apicomplexa Parasites Requires Actin Polymerization in the Host Cell. Cell Host & Microbe 5:259–272.

28. Gilson PR, Crabb BS. 2009. Morphology and kinetics of the three distinct phases of red blood cell invasion by *Plasmodium falciparum* merozoites. International Journal for Parasitology 39:91–96.

29. Weiss GE, Crabb BS, Gilson PR. 2016. Overlaying Molecular and Temporal Aspects of Malaria Parasite Invasion. Trends in Parasitology 32:284–295.

30. Burns AL, Dans MG, Balbin JM, deKoning-Ward TF, Gilson PR, Beeson JG, Boyle MJ, Wilson DW. 2019. Targeting malaria parasite invasion of red blood cells as an antimalarial strategy. doi:10.1093/femsre/fuz005.

31. Van Voorhis WC, Adams JH, Adelfio R, Ahyong V, Akabas MH, Alano P, Alday A, Alemán Resto Y, Alsibaee A, Alzualde A, Andrews KT, Avery SV, Avery VM, Ayong L, Baker M, Baker S, Ben Mamoun C, Bhatia S, Bickle Q, Bounaadja L, Bowling T, Bosch J, Boucher LE, Boyom FF, Brea J, Brennan M, Burton A, Caffrey CR, Camarda G, Carrasquilla M, Carter D, Belen Cassera M, Chih-Chien Cheng K, Chindaudomsate W, Chubb A, Colon BL, Colón-López DD, Corbett Y, Crowther GJ, Cowan N, D’Alessandro S, Le Dang N, Delves M, DeRisi JL, Du AY, Duffy S, Abd El-Salam El-Sayed S, Ferdig MT, Fernández Robledo JA, Fidock DA, et al. 2016. Open Source Drug Discovery with the Malaria Box Compound Collection for Neglected Diseases and Beyond. PLOS Pathogens 12:e1005763.

32. Subramanian G, Belekar MA, Shukla A, Tong JX, Sinha A, Chu TTT, Kulkarni AS, Preiser PR, Reddy DS, Tan KSW, Shanmugam D, Chandramohanadas R. 2018. Targeted Phenotypic Screening in *Plasmodium falciparum* and Toxoplasma gondii Reveals Novel Modes of Action of Medicines for Malaria Venture Malaria Box Molecules. mSphere 3:e00534–17.

33. Azevedo MF, Nie CQ, Elsworth B, Charnaud SC, Sanders PR, Crabb BS, Gilson PR. 2014. *Plasmodium falciparum* Transfected with Ultra Bright NanoLuc Luciferase Offers High Sensitivity Detection for the Screening of Growth and Cellular Trafficking Inhibitors. PLOS ONE 9:e112571.

34. Gurnett AM, Liberator PA, Dulski PM, Salowe SP, Donald RGK, Anderson JW, Wiltsie J, Diaz CA, Harris G, Chang B, Darkin-Rattray SJ, Nare B, Crumley T, Blum PS, Misura AS, Tamas T, Sardana MK, Yuan J, Biftu T, Schmatz DM. 2002. Purification and Molecular Characterization of cGMP-dependent Protein Kinase from Apicomplexan Parasites: A NOVEL CHEMOTHERAPEUTIC TARGET. Journal of Biological Chemistry 277:15913–15922.

35. Boyle MJ, Skidmore M, Dickerman B, Cooper L, Devlin A, Yates E, Horrocks P, Freeman C, Chai W, Beeson JG. 2017. Identification of Heparin Modifications and Polysaccharide Inhibitors of *Plasmodium falciparum* Merozoite Invasion That Have Potential for Novel Drug Development. Antimicrobial Agents and Chemotherapy 61:e00709–17.

36. Zhang J-H, Chung TDY, Oldenburg KR. 1999. A Simple Statistical Parameter for Use in Evaluation and Validation of High Throughput Screening Assays. Journal of Biomolecular Screening 4:67–73.

37. Harris KS, Casey JL, Coley AM, Masciantonio R, Sabo JK, Keizer DW, Lee EF, McMahon A, Norton RS, Anders RF, Foley M. 2005. Binding Hot Spot for Invasion Inhibitory Molecules on *Plasmodium falciparum* Apical Membrane Antigen 1. Infection and Immunity 73:6981–6989.

38. Zhang C, Ondeyka JG, Herath KB, Guan Z, Collado J, Pelaez F, Leavitt PS, Gurnett A, Nare B, Liberator P, Singh SB. 2006. Highly Substituted Terphenyls as Inhibitors of Parasite cGMP-Dependent Protein Kinase Activity. Journal of Natural Products 69:710–712.

39. Green JL, Moon RW, Whalley D, Bowyer PW, Wallace C, Rochani A, Nageshan RK, Howell SA, Grainger M, Jones HM, Ansell KH, Chapman TM, Taylor DL, Osborne SA, Baker DA, Tatu U, Holder AA. 2016. Imidazopyridazine Inhibitors of *Plasmodium falciparum* Calcium-Dependent Protein Kinase 1 Also Target Cyclic GMP-Dependent Protein Kinase and Heat Shock Protein 90 To Kill the Parasite at Different Stages of Intracellular Development. Antimicrobial Agents and Chemotherapy 60:1464–1475.

40. Bansal A, Singh S, More KR, Hans D, Nangalia K, Yogavel M, Sharma A, Chitnis CE. 2013. Characterization of *Plasmodium falciparum* Calcium-dependent Protein Kinase 1 (PfCDPK1) and Its Role in Microneme Secretion during Erythrocyte Invasion. Journal of Biological Chemistry 288:1590–1602.

41. Dennis ASM, Rosling JEO, Lehane AM, Kirk K. 2018. Diverse antimalarials from whole-cell phenotypic screens disrupt malaria parasite ion and volume homeostasis. Scientific Reports 8:8795.

42. Spillman NJ, Kirk K. 2015. The malaria parasite cation ATPase PfATP4 and its role in the mechanism of action of a new arsenal of antimalarial drugs. International journal for parasitology Drugs and drug resistance 5:149–162.

43. Baragaña B, Hallyburton I, Lee MCS, Norcross NR, Grimaldi R, Otto TD, Proto WR, Blagborough AM, Meister S, Wirjanata G, Ruecker A, Upton LM, Abraham TS, Almeida MJ, Pradhan A, Porzelle A, Luksch T, Martínez MS, Luksch T, Bolscher JM, Woodland A, Norval S, Zuccotto F, Thomas J, Simeons F, Stojanovski L, Osuna-Cabello M, Brock PM, Churcher TS, Sala KA, Zakutansky SE, Jiménez-Díaz MB, Sanz LM, Riley J, Basak R, Campbell M, Avery VM, Sauerwein RW, Dechering KJ, Noviyanti R, Campo B, Frearson JA, Angulo-Barturen I, Ferrer-Bazaga S, Gamo FJ, Wyatt PG, Leroy D, Siegl P, Delves MJ, Kyle DE, et al. 2015. A novel multiple-stage antimalarial agent that inhibits protein synthesis. Nature 522:315–320.

44. Ashley EA, Phyo AP. 2018. Drugs in Development for Malaria. Drugs 78:861–879.

45. Baragaña B, Norcross NR, Wilson C, Porzelle A, Hallyburton I, Grimaldi R, Osuna-Cabello M, Norval S, Riley J, Stojanovski L, Simeons FRC, Wyatt PG, Delves MJ, Meister S, Duffy S, Avery VM, Winzeler EA, Sinden RE, Wittlin S, Frearson JA, Gray DW, Fairlamb AH, Waterson D, Campbell SF, Willis P, Read KD, Gilbert IH. 2016. Discovery of a Quinoline-4-carboxamide Derivative with a Novel Mechanism of Action, Multistage Antimalarial Activity, and Potent in Vivo Efficacy. Journal of Medicinal Chemistry 59:9672–9685.

46. Duffy S, Sykes ML, Jones AJ, Shelper TB, Simpson M, Lang R, Poulsen S-A, Sleebs BE, Avery VM. 2017. Screening the Medicines for Malaria Venture Pathogen Box across Multiple Pathogens Reclassifies Starting Points for Open-Source Drug Discovery. Antimicrobial agents and chemotherapy 61:e00379–17.

47. Younis Y, Douelle F, Feng T-S, Cabrera DG, Manach CL, Nchinda AT, Duffy S, White KL, Shackleford DM, Morizzi J, Mannila J, Katneni K, Bhamidipati R, Zabiulla KM, Joseph JT, Bashyam S, Waterson D, Witty MJ, Hardick D, Wittlin S, Avery V, Charman SA, Chibale K. 2012. 3,5-Diaryl-2-aminopyridines as a Novel Class of Orally Active Antimalarials Demonstrating Single Dose Cure in Mice and Clinical Candidate Potential. Journal of Medicinal Chemistry 55:3479–3487.

48. Paquet T, Le Manach C, Cabrera DG, Younis Y, Henrich PP, Abraham TS, Lee MCS, Basak R, Ghidelli-Disse S, Lafuente-Monasterio MJ, Bantscheff M, Ruecker A, Blagborough AM, Zakutansky SE, Zeeman A-M, White KL, Shackleford DM, Mannila J, Morizzi J, Scheurer C, Angulo-Barturen I, Martínez MS, Ferrer S, Sanz LM, Gamo FJ, Reader J, Botha M, Dechering KJ, Sauerwein RW, Tungtaeng A, Vanachayangkul P, Lim CS, Burrows J, Witty MJ, Marsh KC, Bodenreider C, Rochford R, Solapure SM, Jiménez-Díaz MB, Wittlin S, Charman SA, Donini C, Campo B, Birkholtz L-M, Hanson KK, Drewes G, Kocken CHM, Delves MJ, Leroy D, Fidock DA, et al. 2017. Antimalarial efficacy of MMV390048, an inhibitor of *Plasmodium* phosphatidylinositol 4-kinase. Science Translational Medicine 9.

49. Rodríguez F, Rozas I, Kaiser M, Brun R, Nguyen B, Wilson WD, García RN, Dardonville C. 2008. New Bis(2-aminoimidazoline) and Bisguanidine DNA Minor Groove Binders with Potent in Vivo Antitrypanosomal and Antiplasmodial Activity. Journal of Medicinal Chemistry 51:909–923.

50. Le Manach C, Nchinda AT, Paquet T, Gonzàlez Cabrera D, Younis Y, Han Z, Bashyam S, Zabiulla M, Taylor D, Lawrence N, White KL, Charman SA, Waterson D, Witty MJ, Wittlin S, Botha ME, Nondaba SH, Reader J, Birkholtz L-M, Jiménez-Díaz MB, Martínez MS, Ferrer S, Angulo-Barturen I, Meister S, Antonova-Koch Y, Winzeler EA, Street LJ, Chibale K. 2016. Identification of a Potential Antimalarial Drug Candidate from a Series of 2-Aminopyrazines by Optimization of Aqueous Solubility and Potency across the Parasite Life Cycle. Journal of Medicinal Chemistry 59:9890–9905.

51. Boyle MJ, Wilson DW, Richards JS, Riglar DT, Tetteh KKA, Conway DJ, Ralph SA, Baum J, Beeson JG. 2010. Isolation of viable *Plasmodium falciparum* merozoites to define erythrocyte invasion events and advance vaccine and drug development. Proceedings of the National Academy of Sciences 107:14378.

52. Wilson DW, Goodman CD, Sleebs BE, Weiss GE, de Jong NW, Angrisano F, Langer C, Baum J, Crabb BS, Gilson PR, McFadden GI, Beeson JG. 2015. Macrolides rapidly inhibit red blood cell invasion by the human malaria parasite, *Plasmodium falciparum*. BMC biology 13:52–52.

53. Wilson DW, Langer C, Goodman CD, McFadden GI, Beeson JG. 2013. Defining the Timing of Action of Antimalarial Drugs against *Plasmodium falciparum*. Antimicrobial Agents and Chemotherapy 57:1455.

54. Collins CR, Hackett F, Strath M, Penzo M, Withers-Martinez C, Baker DA, Blackman MJ. 2013. Malaria Parasite cGMP-dependent Protein Kinase Regulates Blood Stage Merozoite Secretory Organelle Discharge and Egress. PLOS Pathogens 9:e1003344.

55. Taylor HM, McRobert L, Grainger M, Sicard A, Dluzewski AR, Hopp CS, Holder AA, Baker DA. 2010. The malaria parasite cyclic GMP-dependent protein kinase plays a central role in blood-stage schizogony. Eukaryotic cell 9:37–45.

56. Glushakova S, Humphrey G, Leikina E, Balaban A, Miller J, Zimmerberg J. 2010. New Stages in the Program of Malaria Parasite Egress Imaged in Normal and Sickle Erythrocytes. Current Biology 20:1117–1121.

57. Absalon S, Blomqvist K, Rudlaff RM, DeLano TJ, Pollastri MP, Dvorin JD. 2018. Calcium-Dependent Protein Kinase 5 Is Required for Release of Egress-Specific Organelles in Plasmodium falciparum. mBio 9:e00130–18.

58. Mott BT, Ferreira RS, Simeonov A, Jadhav A, Ang KK-H, Leister W, Shen M, Silveira JT, Doyle PS, Arkin MR, McKerrow JH, Inglese J, Austin CP, Thomas CJ, Shoichet BK, Maloney DJ. 2010. Identification and optimization of inhibitors of Trypanosomal cysteine proteases: cruzain, rhodesain, and TbCatB. Journal of medicinal chemistry 53:52–60.

59. Veale CGL. 2019. Unpacking the Pathogen Box—An Open Source Tool for Fighting Neglected Tropical Disease. ChemMedChem 14:386–453.

60. Glushakova S, Mazar J, Hohmann-Marriott MF, Hama E, Zimmerberg J. 2009. Irreversible effect of cysteine protease inhibitors on the release of malaria parasites from infected erythrocytes. Cellular Microbiology 11:95–105.

61. Koch M, Wright KE, Otto O, Herbig M, Salinas ND, Tolia NH, Satchwell TJ, Guck J, Brooks NJ, Baum J. 2017. *Plasmodium falciparum* erythrocyte-binding antigen 175 triggers a biophysical change in the red blood cell that facilitates invasion. Proceedings of the National Academy of Sciences 114:4225.

62. Tham W-H, Lim NTY, Weiss GE, Lopaticki S, Ansell BRE, Bird M, Lucet I, Dorin-Semblat D, Doerig C, Gilson PR, Crabb BS, Cowman AF. 2015. *Plasmodium falciparum* Adhesins Play an Essential Role in Signalling and Activation of Invasion into Human Erythrocytes. PLOS Pathogens 11:e1005343.

63. Singh S, Alam MM, Pal-Bhowmick I, Brzostowski JA, Chitnis CE. 2010. Distinct external signals trigger sequential release of apical organelles during erythrocyte invasion by malaria parasites. PLoS pathogens 6:e1000746–e1000746.

64. Cowman AF, Berry D, Baum J. 2012. The cellular and molecular basis for malaria parasite invasion of the human red blood cell. The Journal of Cell Biology 198:961–971.

65. Perrin AJ, Collins CR, Russell MRG, Collinson LM, Baker DA, Blackman MJ. 2018. The Actinomyosin Motor Drives Malaria Parasite Red Blood Cell Invasion but Not Egress. mBio 9:e00905–18.

66. Crosnier C, Bustamante LY, Bartholdson SJ, Bei AK, Theron M, Uchikawa M, Mboup S, Ndir O, Kwiatkowski DP, Duraisingh MT, Rayner JC, Wright GJ. 2011. Basigin is a receptor essential for erythrocyte invasion by *Plasmodium falciparum*. Nature 480:534.

67. Chen L, Xu Y, Healer J, Thompson JK, Smith BJ, Lawrence MC, Cowman AF. 2014. Crystal structure of PfRh5, an essential *P. falciparum* ligand for invasion of human erythrocytes. eLife 3:e04187.

68. Yap A, Azevedo MF, Gilson PR, Weiss GE, O’Neill MT, Wilson DW, Crabb BS, Cowman AF. 2014. Conditional expression of apical membrane antigen 1 in *Plasmodium falciparum* shows it is required for erythrocyte invasion by merozoites. Cellular Microbiology 16:642–656.

69. Patel A, Perrin AJ, Flynn HR, Bisson C, Withers-Martinez C, Treeck M, Flueck C, Nicastro G, Martin SR, Ramos A, Gilberger TW, Snijders AP, Blackman MJ, Baker DA. 2019. Cyclic AMP signalling controls key components of malaria parasite host cell invasion machinery. PLOS Biology 17:e3000264.

70. Riglar DT, Richard D, Wilson DW, Boyle MJ, Dekiwadia C, Turnbull L, Angrisano F, Marapana DS, Rogers KL, Whitchurch CB, Beeson JG, Cowman AF, Ralph SA, Baum J. 2011. Super-Resolution Dissection of Coordinated Events during Malaria Parasite Invasion of the Human Erythrocyte. Cell Host & Microbe 9:9–20.

71. Tougan T, Toya Y, Uchihashi K, Horii T. 2019. Application of the automated haematology analyzer XN-30 for discovery and development of anti-malarial drugs. Malaria Journal 18:8.

72. Lu XM, Batugedara G, Lee M, Prudhomme J, Bunnik EM, Le Roch KG. 2017. Nascent RNA sequencing reveals mechanisms of gene regulation in the human malaria parasite *Plasmodium falciparum*. Nucleic acids research 45:7825–7840.

73. Bozdech Z, Llinás M, Pulliam BL, Wong ED, Zhu J, DeRisi JL. 2003. The Transcriptome of the Intraerythrocytic Developmental Cycle of *Plasmodium falciparum*. PLOS Biology 1:e5.

74. Rout S, Mahapatra RK. 2019. In silico analysis of *Plasmodium falciparum* CDPK5 protein through molecular modeling, docking and dynamics. Journal of Theoretical Biology 461:254–267.

75. Pino P, Caldelari R, Mukherjee B, Vahokoski J, Klages N, Maco B, Collins CR, Blackman MJ, Kursula I, Heussler V, Brochet M, Soldati-Favre D. 2017. A multi-stage antimalarial targets the plasmepsins IX and X essential for invasion and egress. Science (New York, NY) 358:522–528.

76. Nasamu AS, Glushakova S, Russo I, Vaupel B, Oksman A, Kim AS, Fremont DH, Tolia N, Beck JR, Meyers MJ, Niles JC, Zimmerberg J, Goldberg DE. 2017. Plasmepsins IX and X are essential and druggable mediators of malaria parasite egress and invasion. Science 358:518.

77. Das S, Hertrich N, Perrin Abigail J, Withers-Martinez C, Collins Christine R, Jones Matthew L, Watermeyer Jean M, Fobes Elmar T, Martin Stephen R, Saibil Helen R, Wright Gavin J, Treeck M, Epp C, Blackman Michael J. 2015. Processing of *Plasmodium falciparum* Merozoite Surface Protein MSP1 Activates a Spectrin-Binding Function Enabling Parasite Egress from RBCs. Cell Host & Microbe 18:433–444.

78. Silmon de Monerri NC, Flynn HR, Campos MG, Hackett F, Koussis K, Withers-Martinez C, Skehel JM, Blackman MJ. 2011. Global Identification of Multiple Substrates for *Plasmodium falciparum* SUB1, an Essential Malarial Processing Protease. Infection and Immunity 79:1086–1097.

79. Baldi DL, Andrews KT, Waller RF, Roos DS, Howard RF, Crabb BS, Cowman AF. 2000. RAP1 controls rhoptry targeting of RAP2 in the malaria parasite *Plasmodium falciparum*. The EMBO journal 19:2435–2443.

80. Howard RF, Narum DL, Blackman M, Thurman J. 1998. Analysis of the processing of *Plasmodium falciparum* rhoptry-associated protein 1 and localization of Pr86 to schizont rhoptries and p67 to free merozoites. Molecular and Biochemical Parasitology 92:111–122.

81. Trager W, Jensen JB. 1976. Human malaria parasites in continuous culture. Science 193:673.

82. Hasenkamp S, Russell KT, Horrocks P. 2012. Comparison of the absolute and relative efficiencies of electroporation-based transfection protocols for *Plasmodium falciparum*. Malaria Journal 11:210.

83. Crick AJ, Tiffert T, Shah SM, Kotar J, Lew VL, Cicuta P. 2013. An Automated Live Imaging Platform for Studying Merozoite Egress-Invasion in Malaria Cultures. Biophysical Journal 104:997–1005.

